# A Luciferase Reporter of Ester Prodrug Delivery to the Cytosol of Intracellular Mycobacteria

**DOI:** 10.64898/2026.11.11.762939

**Authors:** Rachita Dash, Noah Lucas, Andrew Spira, Sarah E. Newkirk, Irene Lepori, Mahendra D. Chordia, Ryan Kenyon, M. Sloan Siegrist, Marcos M. Pires

## Abstract

The bacterial cell envelope restricts access to cytoplasmic targets and is a major reason that otherwise potent antibacterial compounds fail in cells. Masking an anionic carboxylic acid as an ester is a common strategy employed to cross membrane barriers, but it succeeds only if the target cell also hydrolyzes the ester. These two requirements, permeation and cellular unmasking, are rarely measured together in live bacterial cells, and existing reporters cannot confirm the arrival of the unmasked compound into the bacterial cytoplasm. Here we describe a bioluminescent platform that resolves both steps in real time in live mycobacteria using carboxylate esters of D-luciferin. Firefly luciferase, which is expressed in the bacterial cytoplasm, requires that luciferin is unmasked before generating a signal. Across a series of alkyl esters in *Mycobacterium smegmatis*, chain length governed permeation while branching adjacent to the carbonyl governed esterase processing, indicating that the two parameters can be tuned independently. The *n*-butyl ester gave the strongest signal in broth, infected macrophages, and in the pathogen *Mycobacterium avium*, whereas the *tert*-butyl ester was inert in *Mycobacterium avium* and unmasking of the whole series was attenuated at phagosomal pH. Therefore, masking groups are best selected in the organism and environment in which they must act.

## INTRODUCTION

Non-tuberculous mycobacteria (NTM) are environmental organisms found in soil and water that are being increasingly recognized as opportunistic human pathogens. NTM species such as members of the *Mycobacterium avium* complex (MAC) and *Mycobacterium abscessus* cause lung infections, particularly in people with existing lung disease or a weakened immune system. In recognition of their growing public health burden, NTM were recently added to the Watch List of the CDC Antimicrobial Resistance Threats Report in 2026.^1^ NTM infections are difficult to treat, requiring a combination of two to three antibiotics for six months to a year or longer. Despite this clear need for improved therapeutics, few new antimycobacterial agents have reached the clinic in recent decades.^2^ A major contributor to this treatment challenge is the intrinsic resistance conferred by the mycobacterial cell envelope. Because most antibacterial targets reside in the cytoplasm, a compound must cross the full thickness of the cell envelope before it can engage them, so poor intracellular influx is a recurring bottleneck that can limit efficacy regardless of intrinsic potency.^3,4^

The prodrug strategy is a well-established route to cytosolic delivery in drug development.^5–8^ A polar or charged functional group that limits membrane permeation is transiently converted into a neutral, more lipophilic derivative, and the parent molecule is regenerated intracellularly by endogenous chemical or enzymatic processing. The most widespread implementation, employed in both mammalian and microbial systems^9^, masks carboxylic acids as esters. At physiological pH a carboxylic acid is deprotonated, and the resulting anion impedes passive diffusion across lipid membranes; esterification removes that charge, raises lipophilicity, and promotes cellular uptake. Once internalized, endogenous esterases hydrolyze the ester to release the active carboxylic acid *in situ*.

Prodrug delivery is especially challenging in diderm bacteria such as mycobacteria, whose envelope presents an exceptionally formidable permeability barrier.^10–12^ The mycobacterial envelope comprises an outer mycomembrane, an arabinogalactan layer, a peptidoglycan layer, and an inner cytoplasmic membrane.^13^ Compounding this complexity, some mycobacterial species are intracellular pathogens that infect and persist within the phagosomes of host macrophages, a niche that helps them evade host surveillance and clearance. Consequently, prodrug candidates directed at intracellular mycobacteria must traverse the macrophage plasma membrane, the phagosomal membrane, and then the mycobacterial cell envelope to reach the bacterial cytoplasm (**Fig. 1a**). An effective prodrug must therefore satisfy two independent requirements, efficient uptake and unmasking at or before the target site (**Fig. 1b**), and failure at either step is sufficient to compromise activity.

**Figure 1.**
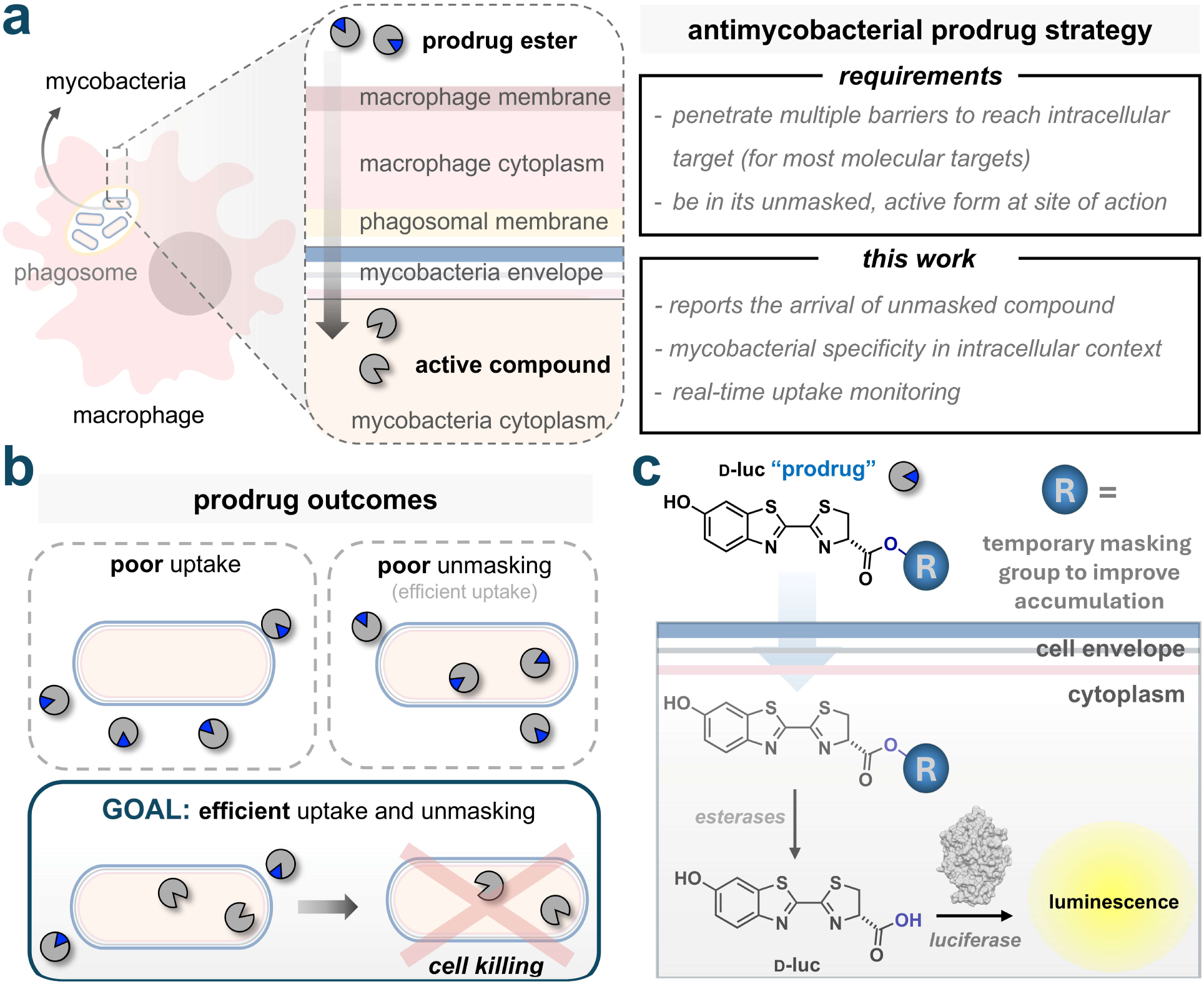
(**a**) The multiple barriers a prodrug must traverse to reach the cytoplasm of mycobacteria residing within host macrophages. (**b**) Possible outcomes of a prodrug strategy; activity requires both efficient cellular uptake and unmasking at the target site. **(c)** The bioluminescence-based assay used to probe arrival of masked esters and its subsequent unmasking in the cytoplasm of mycobacteria.

Here we developed a live-cell, luciferase-based platform that reports both the uptake of ester masking groups into the mycobacterial cytosol and their unmasking within it (**Fig. 1c**). The carboxylic acid of D-luciferin (D-luc) was masked with a series of ester functional groups to generate a panel of model pro-molecules. Because luciferase turns over only the free acid of D-luc, hydrolysis must precede photon emission. Light output is therefore conditional on both cellular uptake and intracellular hydrolysis, which provides a real-time readout of unmasked luciferin in the cytosol of intact cells. Varying the masking group systematically resolved differences among esters within each species and allowed individual esters to be compared across species. We then applied the assay to a macrophage infection model and to the pathogen *Mycobacterium avium* (*Mav*), which showed that pro-moiety performance measured in standard medium was largely conserved in infection-relevant settings, although the most hindered ester of the series fails in the pathogen.

## RESULTS AND DISCUSSION

### Assay development and validation in *Mycobacterium smegmatis*

Ester masking is a potentially viable prodrug strategy for crossing the mycobacterial cell envelope, provided the cell carries esterases capable of unmasking the compound once it arrives. Mycobacteria encode a large complement of esterases and lipases, several with defined roles in lipid metabolism and in entry into or exit from dormancy.^14–19^ Biochemical characterization of individual esterases in live cells has been challenging, in part because many resist heterologous expression and purification.^19–21^ Ester-masked fluorogenic dyes have consequently become the principal tool for surveying this repertoire and have defined global substrate preferences in whole cells and lysates.^22–24^ Bioluminescence offers a complementary and more compartment-restricted observable option. Photons are generated only by substrate turnover, so luciferase expressed in the bacterial cytoplasm stays silent until the free acid of D-luc reaches it. In the case of fluorogenic dye-based assays, a liberated fluorophore, by contrast, can redistribute after hydrolysis and must be excited by incident light, which limits both the depth and the spatial precision of the measurement. A photon is emitted only where enzyme and unmasked substrate meet. Because luciferase-expressing mycobacteria can be detected non-invasively in infected animals,^25^ the same requirement may eventually allow ester processing to be followed during infection.

We first turned to *Escherichia coli* (*E. coli*), as it shares the two-membrane architecture of mycobacteria and because Gram-negative bacteria have themselves been proposed as hosts for prodrug strategies that rely on endogenous hydrolysis.^26^ However, their esterase complements differ from those of mycobacteria, so shared envelope architecture carries no guarantee of shared unmasking behavior. Even in this well-studied organism, findings remain conflicting, with methyl ester hydrolysis reported by us and others,^26–28^ while other work describes minimal activity.^29^ The discrepancy may reflect differences in substrate scope rather than genuine disagreement, since esterase susceptibility depends strongly on the masking group. That variable has received little systematic attention, yet it differs enough between enzymes to determine which organism activates a given prodrug. Resolving the question requires systematic variation of the masking group within a single live-cell assay.

Before deploying a bioluminescent readout, we investigated the presence of bacterial esterases using a conventional fluorescent reporter. Calcein-AM is a nonfluorescent acetoxymethyl ester of calcein, used widely as a mammalian viability stain, where intracellular esterases liberate calcein, an anionic, hydrophilic fluorophore presumed to remain trapped within the cell (**Fig. 2a**).^30,31^ Fluorescence therefore requires both entry of the probe and hydrolysis of its ester groups, which makes it a convenient first comparison of esterase activity between organisms. We therefore incubated *E. coli* and *Mycobacterium smegmatis* (*Msm*), a model organism for mycobacterial pathogens,^32^ with calcein-AM. Fluorescence in *Msm* increased with calcein-AM dosage, whereas *E. coli* remained at background across all concentrations tested (**Fig. 2b**).^33^

**Figure 2.**
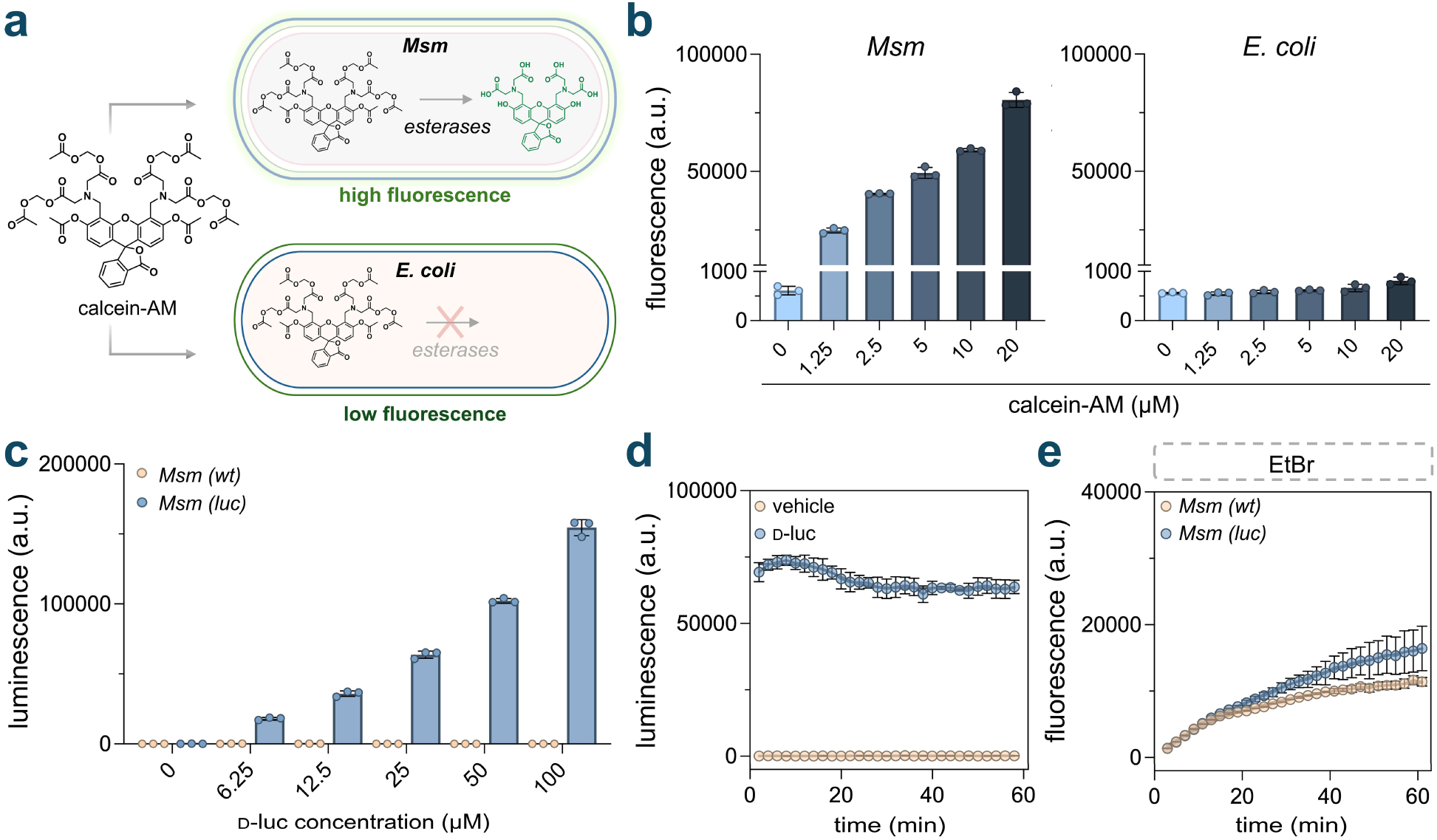
(**a**) Schematic of the calcein-AM assay. (**b**) Fluorescence measured from *E. coli* and *Msm* cells upon addition of varying concentrations of calcein-AM. (**c**) Luminescence of *Msm (luc)* and *Msm (wt)* cells upon incubation with varying concentrations of D-luc at the 60-minute time point. (**d**) Luminescence of *Msm (luc)* after the addition of 25 μM D-luc measured over 60 minutes. (**e**) EtBr-based mycobacterial membrane integrity assay. *Msm (luc)* and *Msm (wt)* cells were incubated with EtBr (5 µM). Fluorescence was monitored for over 60 minutes. Data are represented as mean ± SD (*n* = 3).

The absence of signal in *E. coli* could reflect low esterase expression, inefficient turnover of acetoxymethyl esters, or poor probe accumulation. To distinguish among these possibilities, we monitored calcein-AM fluorescence in *E. coli* ectopically expressing a well-characterized carboxylesterase from *Pseudomonas fluorescens.* At both calcein-AM concentrations tested, esterase-expressing cells showed markedly higher fluorescence than cells lacking the construct, which remained at background (**Fig. S1**). Calcein-AM therefore reaches the *E. coli* cytosol at levels sufficient to generate signal, provided a competent esterase is present. To confirm that the signal arose from esterase activity, we co-treated cells with the serine hydrolase inhibitor diisopropyl fluorophosphate (DIFP). DIFP returned fluorescence to background, indicating that hydrolysis by the ectopically expressed esterase, rather than by endogenous *E. coli* enzymes, drives signal generation. *E. coli* therefore lacks basal esterase activity toward calcein-AM, in contrast to *Msm*.

The calcein-AM results establish two points that frame the next question. Namely, acetoxymethyl esters are processed in *Msm* but not detectably by endogenous *E. coli* enzymes and unmasking inside the cytosol is sufficient to generate signal. What the calcein scaffold cannot report is how broadly mycobacterial esterases act, since the acetoxymethyl group is only one of the pro-moieties used in prodrug design, and calcein-AM interrogates that single chemotype. We therefore moved to a reporter in which the masking group can be varied independently of the scaffold. We posited that esters of D-luc would meet this requirement, since the pro-moiety can be varied across a wide range while the luminophore is held constant, and unmasking inside luciferase-expressing cells produces light in real time with catalytic amplification at the reporting step. We first tuned assay parameters in *Msm* engineered to constitutively express luciferase (*Msm (luc)*) using the parent substrate D-luc. Incubation with varying concentrations of D-luc produced a dose-dependent increase in luminescence relative to wild-type *Msm* (*Msm (wt)*), which gave no appreciable signal at any concentration tested (**Fig. 2c**). These results confirm reporter function and establish that D-luc can access intracellular luciferase in *Msm (luc)* cells. We selected 25 μM as a practical working concentration for D-luc and all its ester derivatives in subsequent assays.

With a working concentration established, we next defined an appropriate incubation window. Time-resolved measurements were expected to report on the relationship between ester structure, uptake, and hydrolysis kinetics, which is informative for prodrug design. We therefore monitored luminescence over time upon addition of 25 μM D-luc to *Msm (luc)* cells (**Fig. 2d**). Luminescence was stable across the 60-minute window. Two features of this profile are worth noting. First, the absence of a measurable lag indicates that permeation of unmodified D-luc across the mycobacterial envelope is fast relative to the sampling interval, so for the parent luminophore the readout is not rate-limited by transport. Second, the plateau indicates that light output is governed by the steady-state concentration of intracellular D-luc rather than by its rate of arrival, consistent with luciferase operating at or near substrate saturation. With the reporting step saturated and transport of the free luminophore fast, differences among the ester derivatives can be attributed to the two steps that precede luciferin release, envelope permeation of the masked compound and esterase-mediated unmasking, rather than to variation in luciferase turnover. We adopted 60 minutes as the standard incubation period for all subsequent assays. Before testing the D-luc ester derivatives, we confirmed that luciferase expression does not itself perturb *Msm* cell envelope integrity (**Fig. 2e**). We used ethidium bromide (EtBr), an established marker of mycobacterial cell envelope integrity.^34–37^ EtBr is weakly fluorescent in aqueous solution and fluoresces strongly upon intercalation into nucleic acids, so its signal in whole cells reflects the extent to which it has crossed the envelope and reached the cytoplasm. Because mycobacteria restrict EtBr entry through the low-permeability mycobacterial cell envelope and extrude it *via* efflux, cells with a compromised envelope accumulate the dye more rapidly and reach higher steady-state fluorescence. If constitutive luciferase expression perturbed envelope assembly or integrity, *Msm* (*luc*) cells should therefore show elevated EtBr fluorescence relative to the parent strain. Instead, *Msm* (*luc*) and *Msm* (*wt*) cells exhibited comparable fluorescence (**Fig. 2e and Fig. S2**), indicating that luciferase expression does not measurably alter envelope permeability and is unlikely to confound the accumulation profiles of the test compounds.

### Behavior of the D-luc ester series in *Mycobacterium smegmatis*

Having established assay parameters, we applied the platform to interrogate mycobacterial esterase activity systematically. We synthesized a focused series of D-luc ester derivatives in which the carboxylate of the luminophore is masked with a simple alkyl group (**Fig. 3a**).^38–40^ The series comprised five members, D-luc-Me, D-luc-Et, D-luc-*i*Pr, D-luc-*n*Bu, and D-luc-*t*Bu, chosen to vary two structural parameters independently. Chain length increases across the linear members, methyl to ethyl to *n*-butyl, raising lipophilicity in a graded manner while leaving the carbonyl sterically accessible. Branching at the α-carbon increases across ethyl, isopropyl, and *tert*-butyl at comparable carbon count, which shields the ester carbonyl from nucleophilic attack with only modest change in lipophilicity. Comparing within each axis therefore separates two effects that are conflated in most prodrug series, namely how a pro-moiety alters partitioning into and across the mycobacterial cell envelope and how it alters the intrinsic susceptibility of the ester bond to enzymatic hydrolysis.

**Figure 3.**
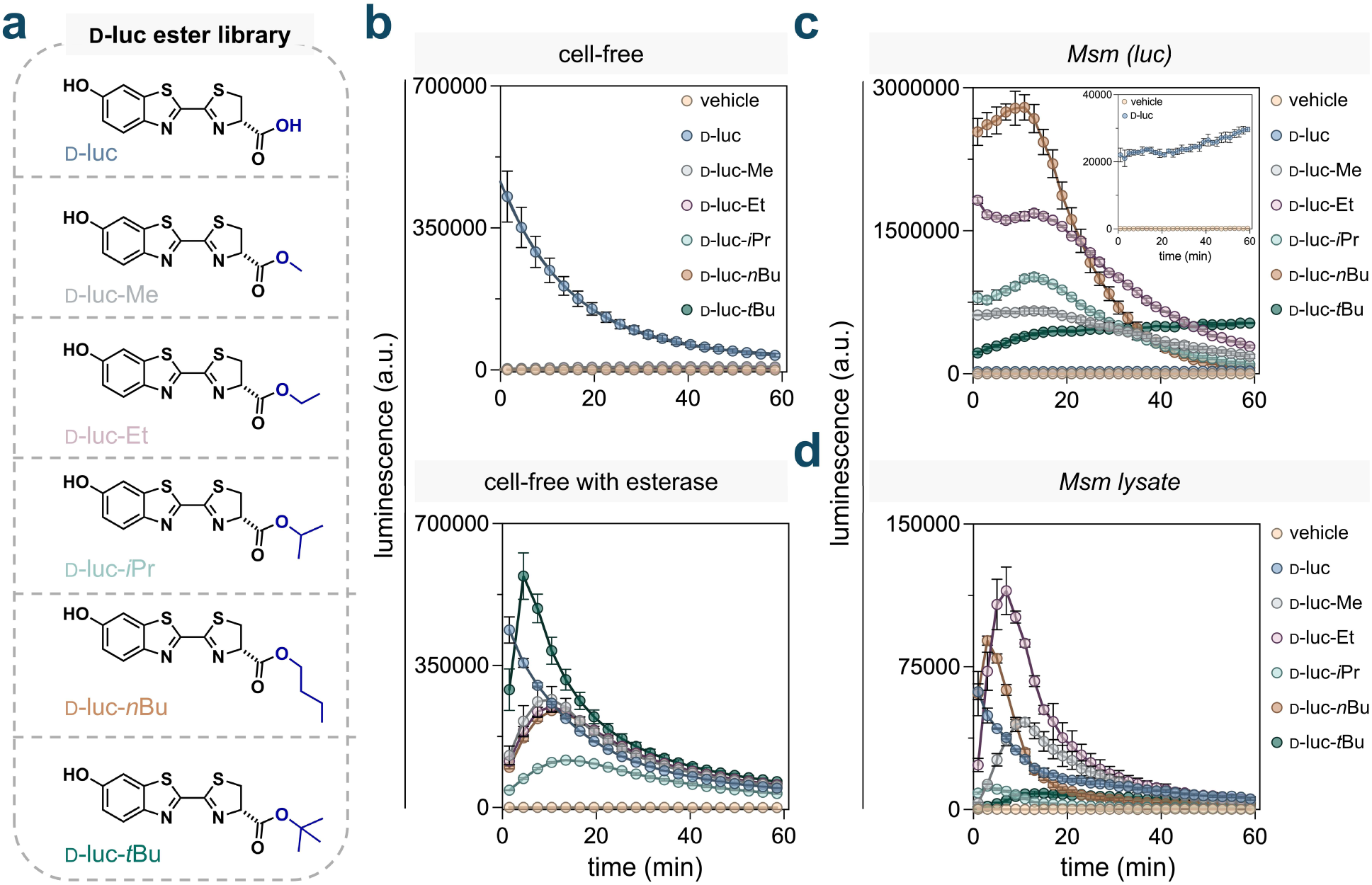
(**a**) Structures of D-luc ester derivatives. (**b**) Cell-free luminescence of luciferase upon the addition of the D-luc ester derivatives, with or without porcine liver esterase (PLE). (**c**) Luminescence of *Msm (luc)* cells after the addition of D-luc ester derivatives measured over 60 minutes. (**d**) Luminescence from cell lysate derived from *Msm (luc)* cells after the addition of D-luc ester derivatives measured over 60 minutes. All compounds were at 25 μM unless otherwise noted. Data are represented as mean ± SD (*n* = 3).

Before evaluating the series in cells, we established the specificity of the reporting step itself. The assay rests on the premise that light is produced only after the ester pro-moiety is removed, which requires that the masked compounds be poor substrates for luciferase. This expectation has a mechanistic basis, because firefly luciferase activates its substrate by adenylation of the C-1 carboxylate and esterification of that position blocks the first chemical step of the reaction.^41^ We therefore measured luminescence from each derivative with purified luciferase in the absence of cells. The parent D-luc gave strong signal, whereas all five esters remained at background (**Fig. 3b**). Direct turnover of the masked compounds is therefore negligible relative to that of free luciferin, and luminescence in the subsequent cellular assays can be assigned to compounds that have both entered the cell and been unmasked.

The preceding result establishes that the esters are not turned over by luciferase, but not that they are competent substrates for hydrolysis. To test this, we supplemented the cell-free system with porcine liver esterase (PLE), a well-characterized carboxylesterase. If the ester pro-moieties are removable, PLE should restore luminescence by generating free D-luc *in situ*, and the magnitude of that recovery should track the susceptibility of each ester to enzymatic hydrolysis. Luminescence increased for all derivatives upon PLE addition, confirming that each member of the series can be unmasked and that the luminophore survives installation and removal of the pro-moiety (**Fig. 3b**). D-luc-*t*Bu yielded the highest signal, despite tertiary esters being the most sterically hindered members of the series. We then asked whether this profile carried over to a more complex esterase milieu by incubating the series in mouse serum, where D-luc-*t*Bu was the only derivative that remained stable over 5 minutes (**Fig. S3**). The inversion indicates that susceptibility to unmasking is not an intrinsic property of the pro-moiety, but a property of the pro-moiety paired with a particular esterase complement. An assay built on a single purified enzyme, however well characterized, is therefore a poor proxy for the hydrolytic environment a prodrug will encounter in the cytosol, and predicting unmasking in a target organism requires measurement in that organism.

With the hydrolytic competence of the series established *in vitro*, we next evaluated its behavior in live *Msm* cells (**Fig. 3c**). Two opposing expectations frame this experiment. Esterification masks the carboxylate, removing an anionic charge that is generally thought to disfavor passive membrane diffusion in mammalian cells.^42–44^ Extending this reasoning to mycobacteria, the esters might be expected to accumulate more efficiently than the parent luminophore and give higher signal on that basis alone. Against this, each ester must be hydrolyzed before light can be produced, and the steric features that improve partitioning (by increasing lipophilicity) may simultaneously impede esterase access. Net luminescence therefore reports the combined outcome of the two steps rather than either one in isolation. All ester-masked derivatives produced considerably higher luminescence than D-luc itself (**Fig. 3c**, **inset**), consistent with charge neutralization as the dominant determinant of entry. This also means that any unintended hydrolysis in the extracellular space would contribute little signal, since the liberated parent accumulates poorly. Within the series, D-luc-*n*Bu gave the highest signal and responded in a concentration-dependent manner (**Fig. S4**). The branching axis resolved in the opposite direction: D-luc-Et exceeded D-luc-*i*Pr, and D-luc-*t*Bu gave substantially lower signal than D-luc-*n*Bu despite carrying the same number of carbons. That last pair is the most informative comparison in the dataset, since the two derivatives are matched in carbon count and closely matched in lipophilicity, leaving steric access to the carbonyl as the principal variable that distinguishes them. Branching near the ester carbonyl, rather than chain length or lipophilicity, therefore, appears to limit unmasking in *Msm*, a conclusion we test directly below. Treatment with the D-luc ester series at the tested concentrations did not perturb *Msm (luc)* cells, as assessed by the EtBr assay (**Fig. S5**).

Taken together, the cellular data report the combined outcome of permeation and hydrolysis, so the ordering among derivatives cannot by itself be assigned to either step. To separate them, we repeated the measurement in lysates prepared from *Msm* (*luc*) (**Fig. 3d**). Disrupting the cell envelope removes the physical barrier to cytosolic entry while retaining the endogenous esterase complement and luciferase, so differences that persist in lysate are attributable to substrate preference at the hydrolysis step, while differences that collapse point to uptake-based factors. We expected free D-luc to give the strongest and fastest signal in this format, since it bypasses the unmasking requirement entirely, but it did not. This discrepancy is likely a property of the lysate system rather than of the substrate. Luciferase turnover consumes ATP in the adenylation step, and in live cells ATP is continuously regenerated so that the enzyme can cycle. A lysate may not possess such regeneration capacity, so the available ATP pool supports what amounts to a single round of turnover. Free D-luc, requiring no prior processing, likely saturates that pool immediately upon addition and is consumed in a burst that precedes our first measurement. Consistent with this, D-luc was the only substrate for which no initial rise was captured, so its trace likely represents the tail of a reaction that had already largely run to completion.

Within that constraint, the lysate format is well suited to comparing the pro-moieties against one another, since all five are released gradually by esterase action and draw on the ATP pool across the measurement window, which places them under identical conditions. Reading the two formats against each other resolves the series into contributions that neither measurement provides on its own. A pro-moiety that performs well in both is efficiently taken up and efficiently hydrolyzed. One that performs well in cells but falls back in lysate owes its cellular advantage to permeation. One that performs poorly in both is limited at the hydrolysis step, since removing the envelope does nothing to rescue it. D-luc-*n*Bu yielded the strongest signal in intact cells but was overtaken by D-luc-Et in lysate. Because both are unbranched and present the same unhindered carbonyl to esterases, they should be comparably good substrates, and the lysate result confirms that they are, with D-luc-Et modestly ahead. The higher signal for D-luc-*n*Bu in whole cells therefore reflects greater accumulation, consistent with its longer and more lipophilic chain promoting accumulation. This is the clearest demonstration in the dataset that the two steps can be separated. The same pair of compounds reverses order depending on whether the envelope is present, and the direction of the reversal identifies which step dominates each format. The branched esters behaved differently. D-luc-*i*Pr trailed D-luc-Et in both intact cells and lysate, and D-luc-*t*Bu remained at the bottom of the series in both formats despite matching D-luc-*n*Bu in carbon count and closely matching it in lipophilicity. Because the envelope is absent in lysate, these deficits must arise at the hydrolysis step, which establishes that branching impairs mycobacterial ester processing. Chain length therefore modulates accumulation, with an effect seen primarily in intact cells, whereas branching modulates hydrolysis, with an effect that persists wherever esterases are present. This separation matters for prodrug design in mycobacteria, because the two parameters can be tuned independently. Lipophilicity can be raised to improve entry/retention without compromising unmasking, provided the added carbon is not placed adjacent to the carbonyl. These trends are consistent with prior studies showing that mycobacterial esterases preferentially hydrolyze longer chain ester substrates, including alkyl chains of four carbons.^19,21,45–48^

We next applied the same analysis to *E. coli* expressing luciferase (*E. coli* (*luc*)), where the calcein-AM experiments had already indicated a deficit in endogenous esterase activity. Two outcomes were possible. If *E. coli* possesses esterases competent for these pro-moieties, the pattern should resemble that in *Msm*, with the masked derivatives outperforming the charged parent on the strength of improved accumulation. If it does not, the esters should perform worse than D-luc regardless of how efficiently they accumulate, since unmasking is required before light can be produced and no amount of accumulation can compensate for its absence. All ester-masked derivatives gave lower signals than parent D-luc, with D-luc-Me highest among them (**Fig. S6**). On its own this result is ambiguous, since the assay cannot distinguish a compound that never entered from one that entered and was never unmasked. To resolve this, we generated a strain co-expressing luciferase and the carboxylesterase from *Pseudomonas fluorescens*, as we had done for the calcein-AM experiment. If the esters reach the cytosol but lack a native enzyme to unmask them, supplying a competent esterase should increase signal output; if they do not reach the cytosol, signal should remain low. Ectopic expression of the esterase in *E. coli* (*luc*) raised D-luc-Me and D-luc-Et above parent D-luc, while the remaining three esters stayed below it (**Fig. S6**). The methyl and ethyl esters therefore reach the *E. coli* cytosol at functionally relevant levels, and their weak signals in the parent strain reflect the absence of endogenous esterases that recognize them rather than an accumulation barrier. The larger esters remained low even with the exogenous esterase present, consistent with poor recognition by that enzyme and paralleling the steric trend established in *Msm*. The two organisms thus fail to unmask these pro-moieties for different reasons. In *Msm* the limitation is structural and confined to branched esters that resist an otherwise capable esterase complement, whereas in *E. coli* the limitation is the absence of hydrolytic activity itself, which leaves even the smallest and most accessible esters intact. Ester masking is therefore not a general strategy for delivering charged cargo into bacteria; it works only where the target organism supplies the enzyme to undo it.

### Behavior of the D-luc ester series in infection-relevant contexts

The measurements to this point were made at neutral pH, which does not represent the environment in which mycobacteria are encountered during infection. Pathogenic mycobacteria reside and replicate within macrophage phagosomes, a compartment substantially more acidic than either the bacterial or the host cytosol, and a pro-moiety optimized under neutral conditions may behave differently there. Esterase catalysis depends on the protonation state of the active-site serine-histidine pair, so hydrolysis rates are pH-dependent. The bacterium may also adjust its physiology under acid stress, which might change the abundance of the esterases available to act.^49,50^ Because our readout integrates accumulation and unmasking, these effects are not separable within a single measurement, but their net consequence for prodrug activation is the quantity of practical interest. We therefore incubated the D-luc derivatives with *Msm* (*luc*) at reduced pH (**Fig. 4a**). All ester derivatives produced diminished luminescence relative to neutral pH within the kinetic window of the assay. The direction of this effect matches a prior report that esterase activity in *Msm* homogenate toward ethyl benzoate declines as pH is lowered from 7.0 to 5.5,^51^ which points to the hydrolysis step rather than entry as the dominant contribution. Pro-moiety performance is therefore not a fixed property of the compound: a masking group selected based on activation in broth may be activated far less efficiently in the compartment where the bacterium resides, a variable that assays conducted at neutral pH cannot reveal.

**Figure 4.**
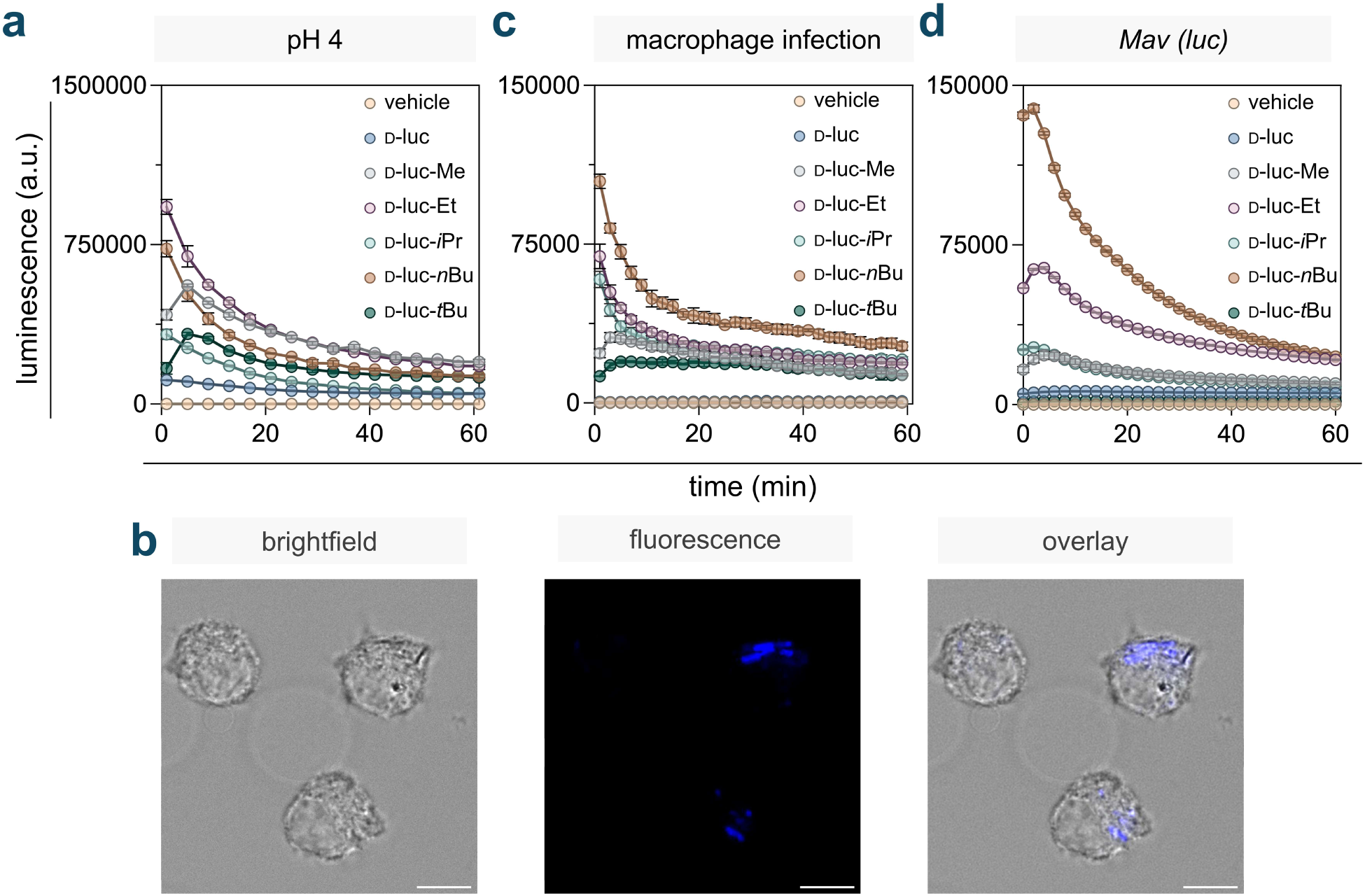
(**a**) Luminescence of *Msm (luc)* cells after the addition of D-luc ester derivatives measured over 60 minutes at pH 4. (**b**) Confocal analysis of macrophages infected with DADA-labeled (100 µM) *Msm (wt)*; scale bar = 10 µm. (**c**) Luminescence of macrophages infected with *Msm (luc)* cells after the addition of D-luc ester derivatives measured over 60 minutes. (**d**) Luminescence of *Mav (luc)* cells after the addition of D-luc ester derivatives measured over 60 minutes. All compounds in the luciferase assays were at 25 μM unless otherwise noted. Data are represented as mean ± SD (*n* = 3).

Acidification also reordered the ester series relative to neutral pH. D-luc-Et yielded the highest signal among the esters at pH 4, whereas D-luc-*n*Bu was highest at pH 7, indicating that the relative merit of a pro-moiety depends on the compartment in which it is assessed. As in the lysate experiment, part of the apparent loss of signal at low pH may be a limitation of temporal resolution. If acidification accelerates the early phase of the reaction, a rapid luminescence burst following substrate addition would fall outside our first measurement, and the integrated signal would understate what was produced. The parent compound bears directly on this concern. D-luc displayed substantially elevated signal at low pH (**Fig. S7**), consistent with improved accumulation of the protonated carboxylic acid, more efficient luciferase turnover under acidic conditions, or both. Because the parent requires no unmasking, its behavior isolates the pH dependence of the accumulation and reporting steps from that of hydrolysis. That the esters move in the opposite direction while the parent gains signal argues that the decline reflects impaired esterase activity rather than a general suppression of the readout, in agreement with the homogenate measurement described above.

Acidification of the medium reproduces one feature of the phagosome but not the full route an antimycobacterial compound must take to reach the cytosol of the pathogen. In an infected host cell, a molecule must cross the macrophage plasma membrane and then the phagosomal membrane before it encounters the bacterial envelope, and the route it takes can change which pro-moiety is appropriate. A compound that partitions directly into the macrophage cytosol is exposed to mammalian carboxylesterases, which are abundant and act readily on these pro-moieties. Unmasking in the mammalian cytosol before arrival at the pathogen liberates a charged molecule, which we expect to be poorly positioned to cross the phagosomal membrane and reach the bacterium. The two routes impose opposite requirements on a pro-moiety, and no bacteria-only measurement can distinguish them, since broth contains no host compartment in which a compound can be lost before it reaches its target. Resolving this experimentally has been difficult because standard reporters cannot localize the event they detect. A fluorogenic ester taken up by an infected macrophage could be unmasked by either host cytosolic esterases or by bacterial ones, and the resulting fluorophore reports a hydrolysis that may have occurred anywhere in the cell. The two-component design of the D-luc system supplies the missing spatial constraint. Luciferase is expressed only in the bacterium, so photon production requires that the unmasked luminophore occupy the mycobacterial cytosol regardless of where hydrolysis took place. Free D-luc, bearing an anionic carboxylate, enters mycobacteria poorly and generates minimal signal (**Fig. 3c**), so luminophore liberated prematurely in the host cytosol is largely excluded from the bacterium. We therefore adapted the assay to a macrophage infection model, which to our knowledge provides the first direct readout of pro-moiety unmasking inside intracellular mycobacteria, measured in the compartment where the reaction matters rather than inferred from bacteria in broth.

Establishing the model required first confirming that the bacteria were intracellular. We metabolically labeled the peptidoglycan of *Msm* using Diethyl-Amino-coumarin-D-Alanine (DADA), a fluorescent D-amino acid that marks the envelope of living cells.^52–55^ Labeled cells were readily visualized within J774A.1 macrophages by confocal microscopy, confirming internalization (**Fig. 4b**). We then incubated infected macrophages with the D-luc ester series (**Fig. 4c**). If an ester reaches the bacterial cytoplasm with its pro-moiety intact, it should generate signal comparable to that obtained with isolated bacteria. Ester-masked derivatives generated substantially higher luminescence than unmasked D-luc, with an initial spike indicating rapid entry and processing. Relative performance across the series closely paralleled the bacteria-only measurements. Two features of this result are notable. First, the magnitude of the ester advantage over parent D-luc indicates that a substantial fraction of each compound traverses the host membranes with its pro-moiety intact, since extensive premature hydrolysis in the macrophage cytosol would have compressed that difference. Second, preservation of the rank order across the series indicates that the determinants identified in broth, cell accumulation and steric access to the ester carbonyl, remain the controlling variables once the bacterium is inside a host cell. Pro-moiety performance measured in broth therefore translates to the intracellular setting for this series, and the host compartment does not impose a separate selection on which masking groups succeed.

The determinants established so far come from *Msm*, a fast-growing environmental organism chosen for tractability rather than for clinical relevance. Whether these pro-moiety trends extend to a pathogen require direct testing. We therefore evaluated the D-luc ester series in *Mycobacterium avium* (*Mav*), a slow-growing non-tuberculous mycobacterium that causes pulmonary disease, particularly in immunocompromised individuals. *Mav* was engineered to express luciferase constitutively (*Mav* (*luc*)) and assayed under conditions matched to those used for *Msm* (*luc*) (**Fig. 4d**). The dominant trend was preserved. As in *Msm*, D-luc-nBu produced a markedly stronger signal than the other derivatives, indicating that the balance of lipophilicity and steric accessibility that favors the linear butyl ester holds across both species. The series diverged at its most hindered member. D-luc-*t*Bu gave no appreciable signal in *Mav*, whereas it remained detectable in *Msm.* The assay cannot distinguish whether this reflects an esterase complement unable to act on a tertiary ester or poor accumulation of that compound in *Mav*, and the two possibilities are not mutually exclusive. What the result does establish is that the steric penalty identified in *Msm* is not uniform across mycobacteria but becomes more severe in a pathogenic species, such that a pro-moiety retaining partial activity in the model organism can be entirely inert in the intended target. For prodrug design, a masking group should therefore be validated in the species of interest rather than extrapolated from *Msm*, and the platform described here provides a direct means of doing so in any mycobacterium that tolerates luciferase expression.

## CONCLUSION

We developed a bioluminescence-based platform that profiles esterase preferences in live mycobacteria using carboxylate ester derivatives of D-luc as reporter substrates. Its advantage over existing reporters is that the readout is conditional on cytosolic arrival of the unmasked acid within the bacterium itself, because luciferase is confined to the bacterial cytoplasm and turns over only the free carboxylate. Light therefore reports productive delivery rather than hydrolysis at an unspecified location, which is the quantity that matters for prodrugs directed at intracellular targets.

Applying the platform to a focused series of D-luc esters varying in alkyl chain length and steric bulk, we found that ester masking consistently enhanced luminescent output relative to free D-luc, consistent with improved cytosolic accumulation. Comparing intact cells with lysates then separated the two steps that the cellular readout combines. Chain length acted mainly on accumulation, since the advantage of D-luc-*n*Bu over D-luc-Et in whole cells disappeared once the envelope was removed. Branching adjacent to the carbonyl acted on hydrolysis, since the deficits of D-luc-*i*Pr and D-luc-*t*Bu persisted in lysate. D-luc-*n*Bu produced the strongest signal overall, identifying the *n*-butyl ester as a promising masking group for prodrug applications, and the independence of the two parameters means that lipophilicity can be raised to improve entry without sacrificing unmasking.

Delivery to mycobacteria is further complicated by their intracellular life cycle. During infection, mycobacterial pathogens can reside within phagosomal compartments of host macrophages.^56^ Compounds must therefore cross both the macrophage plasma membrane and the phagosomal membrane before reaching the bacterial cell envelope. Adapting the assay to a macrophage incubation model showed that the rank order established in broth is preserved inside host cells, indicating that a substantial fraction of each ester crosses the host membranes with its pro-moiety intact. Extending the assay to *Mav*, a slow-growing pathogenic species, showed that the trends largely generalize, with the notable exception of the *tert*-butyl ester, which was inert in *Mav* while remaining detectable in *Msm*. Acidification to phagosomal pH attenuated unmasking across the series and reordered it, so neither the species nor the compartment can be assumed from a measurement made in broth at neutral pH.

Two limitations bound these conclusions. The readout integrates accumulation and hydrolysis, so assigning an effect to a single step requires the paired lysate comparison rather than the cellular measurement alone, and the kinetic window may under-sample rapid bursts, a caveat that applies most directly to the lysate and low-pH datasets. Further, the *Mav* result shows that extrapolation between mycobacterial species is not safe. With those caveats, this work establishes a foundation for the rational design of ester-based prodrugs targeting intracellular mycobacteria, leveraging the endogenous esterase activity of the mycobacterial cytosol as a trigger. The structural determinants of ester uptake and hydrolysis defined here, together with the platform itself, should transfer directly to prodrug development aimed at improving intracellular delivery of antibacterial agents against pathogenic mycobacteria, including *non-tuberculous mycobacteria like Mav*.

## ACKNOWLEDGMENTS

This work was supported by R35GM124893, R01AI191591-01, R01AI179080-01, and R01AI200342-01. We thank the director of the University of Massachusetts Amherst Biophysical Characterization facilities, Dr. Stephen Eyles, for his help and advice, and the Massachusetts Life Science Center for the funding for the equipment.

## SUPPORTING INFORMATION

Additional figures, tables, and materials and methods are provided in the Supporting Information file.

## REFERENCES

(1) Antimicrobial Resistance Threats | AR Threats Report. https://arpsp.cdc.gov/artr/antimicrobial-resistance-threats (accessed 2026-11-11)

(2) Farhat, M.; Cox, H.; Ghanem, M.; Denkinger, C. M.; Rodrigues, C.; Abd El Aziz, M. S.; Enkh-Amgalan, H.; Vambe, D.; Ugarte-Gil, C.; Furin, J.; Pai, M. Drug-Resistant Tuberculosis: A Persistent Global Health Concern. Nat. Rev. Microbiol. 2024, 22 (10), 617–635. 10.1038/s41579-024-01025-1

(3) Silver, L. L. Challenges of Antibacterial Discovery. Clin. Microbiol. Rev. 2011, 24 (1), 71–109. 10.1128/cmr.00030-10

(4) Tommasi, R.; Brown, D. G.; Walkup, G. K.; Manchester, J. I.; Miller, A. A. ESKAPEing the Labyrinth of Antibacterial Discovery. Nat. Rev. Drug Discov. 2015, 14 (8), 529– 542. 10.1038/nrd4572

(5) Taylor, M. D. Improved Passive Oral Drug Delivery via Prodrugs. Adv. Drug Deliv. Rev. 1996, 19 (2), 131–148. 10.1016/0169-409X(95)00104-F

(6) Beaumont, K.; Webster, R.; Gardner, I.; Dack, K. Design of Ester Prodrugs to Enhance Oral Absorption of Poorly Permeable Compounds: Challenges to the Discovery Scientist. http://www.eurekaselect.com 2003, 4 (6).

(7) Ettmayer, P.; Amidon, G. L.; Clement, B.; Testa, B. Lessons Learned from Marketed and Investigational Prodrugs. J. Med. Chem. 2004, 47 (10), 2393–2404. 10.1021/jm0303812

(8) Rautio, J.; Kumpulainen, H.; Heimbach, T.; Oliyai, R.; Oh, D.; Järvinen, T.; Savolainen, J. Prodrugs: Design and Clinical Applications. Nat. Rev. Drug Discov. 2008, 7 (3), 255–270. 10.1038/nrd2468

(9) Hetrick, K. J.; Raines, R. T. Assessing and Utilizing Esterase Specificity in Antimicrobial Prodrug Development. Methods Enzymol. 2022, 664, 199–220. 10.1016/bs.mie.2021.11.008

(10) Lepori, I.; Liu, Z.; Evbarunegbe, N.; Feng, S.; Brown, T. P.; Mane, K.; Dash, R.; Shivangi; Wong, M.; Naick, A.; George, A.; Guo, T.; Shelke, A. M.; Sharpless, K. B.; Dong, J.; Freundlich, J. S.; Im, W.; Green, A. G.; Pires, M. M.; Siegrist, M. S. Identification of Chemical Features for Improved Outer Membrane Permeation in Mycobacteria Using Machine Learning. Nat. Microbiol. 2026, 11 (9), 2647–2665. 10.1038/s41564-026-02412-5

(11) Lepori, I.; Jackson, K.; Liu, Z.; Chordia, M. D.; Wong, M.; Rivera, S. L.; Roncetti, M.; Poliseno, L.; Freundlich, J. S.; Pires, M. M.; Siegrist, M. S. The Mycomembrane Differentially and Heterogeneously Restricts Antibiotic Permeation. ACS Infect. Dis. 2025, 11 (7), 1893–1906. 10.1021/acsinfecdis.4c01062

(12) Dash, R.; Liu, Z.; Lepori, I.; Chordia, M. D.; Ocius, K.; Holsinger, K.; Zhang, H.; Kenyon, R.; Im, W.; Siegrist, M. S.; Pires, M. M. Systematic Determination of the Impact of Structural Edits on Peptide Accumulation into Mycobacteria. ACS Chem. Biol. 2025, 20 (8), 1962–1979. 10.1021/acschembio.5c00330

(13) Daffé, M.; Marrakchi, H. Unraveling the Structure of the Mycobacterial Envelope. Microbiol. Spectr. 2019, 7 (4), 10.1128/microbiolspec.gpp3-0027-2018. https://doi.org/10.1128/microbiolspec.gpp3-0027-2018

(14) Dang, G.; Chen, L.; Li, Z.; Deng, X.; Cui, Y.; Cao, J.; Yu, S.; Pang, H.; Liu, S. Expression, Purification and Characterisation of Secreted Esterase Rv2525c from Mycobacterium Tuberculosis. Appl. Biochem. Biotechnol. 2015, 176 (1), 1–12. 10.1007/s12010-015-1555-9

(15) Chen, L.; Dang, G.; Deng, X.; Cao, J.; Yu, S.; Wu, D.; Pang, H.; Liu, S. Characterization of a Novel Exported Esterase Rv3036c from *Mycobacterium Tuberculosis*. Protein Expr. Purif. 2014, 104, 50–56. 10.1016/j.pep.2014.09.003

(16) Sultana, R.; Vemula, M. H.; Banerjee, S.; Guruprasad, L. The PE16 (Rv1430) of Mycobacterium Tuberculosis Is an Esterase Belonging to Serine Hydrolase Superfamily of Proteins. PLOS ONE 2013, 8 (2), e55320. 10.1371/journal.pone.0055320

(17) Lun, S.; Bishai, W. R. Characterization of a Novel Cell Wall-Anchored Protein with Carboxylesterase Activity Required for Virulence in *Mycobacterium Tuberculosis*\*. J. Biol. Chem. 2007, 282 (25), 18348–18356. 10.1074/jbc.M700035200

(18) Singh, K. H.; Jha, B.; Dwivedy, A.; Choudhary, E.; N, A. G.; Ashraf, A.; Arora, D.; Agarwal, N.; Biswal, B. K. Characterization of a Secretory Hydrolase from *Mycobacterium Tuberculosis* Sheds Critical Insight into Host Lipid Utilization by *M. Tuberculosis*. J. Biol. Chem. 2017, 292 (27), 11326–11335. 10.1074/jbc.M117.794297

(19) West, N. P.; Chow, F. M. E.; Randall, E. J.; Wu, J.; Chen, J.; Ribeiro, J. M. C.; Britton, W. J. Cutinase-like Proteins of Mycobacterium Tuberculosis: Characterization of Their Variable Enzymatic Functions and Active Site Identification. FASEB J. 2009, 23 (6), 1694–1704. 10.1096/fj.08-114421

(20) Deb, C.; Daniel, J.; Sirakova, T. D.; Abomoelak, B.; Dubey, V. S.; Kolattukudy, P. E. A Novel Lipase Belonging to the Hormone-Sensitive Lipase Family Induced under Starvation to Utilize Stored Triacylglycerol in Mycobacterium Tuberculosis. J. Biol. Chem. 2006, 281 (7), 3866–3875. 10.1074/jbc.M505556200

(21) Delorme, V.; Diomandé, S. V.; Dedieu, L.; Cavalier, J.-F.; Carrière, F.; Kremer, L.; Leclaire, J.; Fotiadu, F.; Canaan, S. MmPPOX Inhibits Mycobacterium Tuberculosis Lipolytic Enzymes Belonging to the Hormone-Sensitive Lipase Family and Alters Mycobacterial Growth. PLOS ONE 2012, 7 (9), e46493. 10.1371/journal.pone.0046493

(22) Tallman, K. R.; Levine, S. R.; Beatty, K. E. Small-Molecule Probes Reveal Esterases with Persistent Activity in Dormant and Reactivating Mycobacterium Tuberculosis. ACS Infect. Dis. 2016, 2 (12), 936–944. 10.1021/acsinfecdis.6b00135

(23) Tallman, K. R.; Levine, S. R.; Beatty, K. E. Profiling Esterases in Mycobacterium Tuberculosis Using Far-Red Fluorogenic Substrates. ACS Chem. Biol. 2016, 11 (7), 1810–1815. 10.1021/acschembio.6b00233

(24) Bassett, B.; Waibel, B.; White, A.; Hansen, H.; Stephens, D.; Koelper, A.; Larsen, E. M.; Kim, C.; Glanzer, A.; Lavis, L. D.; Hoops, G. C.; Johnson, R. J. Measuring the Global Substrate Specificity of Mycobacterial Serine Hydrolases Using a Library of Fluorogenic Ester Substrates. ACS Infect. Dis. 2018, 4 (6), 904–911. 10.1021/acsinfecdis.7b00263

(25) Andreu, N.; Zelmer, A.; Fletcher, T.; Elkington, P. T.; Ward, T. H.; Ripoll, J.; Parish, T.; Bancroft, G. J.; Schaible, U.; Robertson, B. D.; Wiles, S. Optimisation of Bioluminescent Reporters for Use with Mycobacteria. PLOS ONE 2010, 5 (5), e10777. 10.1371/journal.pone.0010777

(26) Hetrick, K. J.; Aguilar Ramos, M. A.; Raines, R. T. Endogenous Enzymes Enable Antimicrobial Activity. ACS Chem. Biol. 2021, 16 (5), 800–805. 10.1021/acschembio.0c00894

(27) Brown, A. R.; Wodzanowski, K. A.; Santiago, C. C.; Hyland, S. N.; Follmar, J. L.; Asare-Okai, P.; Grimes, C. L. Protected N-Acetyl Muramic Acid Probes Improve Bacterial Peptidoglycan Incorporation via Metabolic Labeling. ACS Chem. Biol. 2021, 16 (10), 1908–1916. 10.1021/acschembio.1c00268

(28) Dash, R.; Holsinger, K. A.; Chordia, M. D.; Gh., M. S.; Pires, M. M. Bioluminescence-Based Determination of Cytosolic Accumulation of Antibiotics in Escherichia Coli. ACS Infect. Dis. 2024, 10 (5), 1602–1611. 10.1021/acsinfecdis.3c00684

(29) Antonczak, A. K.; Simova, Z.; Tippmann, E. M. A Critical Examination of Escherichia Coli Esterase Activity*. J. Biol. Chem. 2009, 284 (42), 28795–28800. 10.1074/jbc.M109.027409

(30) Papadopoulos, N. G.; Dedoussis, G. V. Z.; Spanakos, G.; Gritzapis, A. D.; Baxevanis, C. N.; Papamichail, M. An Improved Fluorescence Assay for the Determination of Lymphocyte-Mediated Cytotoxicity Using Flow Cytometry. J. Immunol. Methods 1994, 177 (1), 101–111. 10.1016/0022-1759(94)90147-3

(31) Lichtenfels, R.; Biddison, W. E.; Schulz, H.; Vogt, A. B.; Martin, R. CARE-LASS (Calcein-Release-Assay), an Improved Fluorescence-Based Test System to Measure Cytotoxic T Lymphocyte Activity. J. Immunol. Methods 1994, 172 (2), 227–239. 10.1016/0022-1759(94)90110-4

(32) Sparks, I. L.; Derbyshire, K. M.; Jacobs, W. R.; Morita, Y. S. Mycobacterium Smegmatis: The Vanguard of Mycobacterial Research. J. Bacteriol. 2023, 205 (1), e00337–22. 10.1128/jb.00337-22

(33) Rego, E. H.; Audette, R. E.; Rubin, E. J. Deletion of a Mycobacterial Divisome Factor Collapses Single-Cell Phenotypic Heterogeneity. Nature 2017, 546 (7656), 153–157. 10.1038/nature22361

(34) Bisson, G. P.; Mehaffy, C.; Broeckling, C.; Prenni, J.; Rifat, D.; Lun, D. S.; Burgos, M.; Weissman, D.; Karakousis, P. C.; Dobos, K. Upregulation of the Phthiocerol Dimycocerosate Biosynthetic Pathway by Rifampin-Resistant, rpoB Mutant Mycobacterium Tuberculosis. J. Bacteriol. 2012, 194 (23), 6441–6452. 10.1128/jb.01013-12

(35) Campodónico, V. L.; Rifat, D.; Chuang, Y.-M.; Ioerger, T. R.; Karakousis, P. C. Altered Mycobacterium Tuberculosis Cell Wall Metabolism and Physiology Associated With RpoB Mutation H526D. Front. Microbiol. 2018, 9. 10.3389/fmicb.2018.00494

(36) Papadopoulos, A. O.; Ealand, C.; Gordhan, B. G.; VanNieuwenhze, M.; Kana, B. D. Characterisation of a Putative M23-Domain Containing Protein in Mycobacterium Tuberculosis. PLOS ONE 2021, 16 (11), e0259181. 10.1371/journal.pone.0259181

(37) Rodrigues, L.; Ramos, J.; Couto, I.; Amaral, L.; Viveiros, M. Ethidium Bromide Transport across Mycobacterium Smegmatis Cell-Wall: Correlation with Antibiotic Resistance. BMC Microbiol. 2011, 11 (1), 35. 10.1186/1471-2180-11-35

(38) Ogasa, C.; Kayano, K.; Namba, K. A Simple and Powerful Tert-Butylation of Carboxylic Acids and Alcohols. Synlett 2024, 35 (2), 235–239. 10.1055/a-2161-9689

(39) Rothweiler, U.; Eriksson, J.; Stensen, W.; Leeson, F.; Engh, R. A.; Svendsen, J. S. Luciferin and Derivatives as a DYRK Selective Scaffold for the Design of Protein Kinase Inhibitors. Eur. J. Med. Chem. 2015, 94, 140–148. 10.1016/j.ejmech.2015.02.035

(40) Craig, F. F.; Simmonds, A. C.; Watmore, D.; McCapra, F.; White, M. R. H. Membrane-Permeable Luciferin Esters for Assay of Firefly Luciferase in Live Intact Cells. Biochem. J. 1991, 276 (3), 637–641. 10.1042/bj2760637

(41) Kaskova, Z. M.; Tsarkova, A. S.; Yampolsky, I. V. 1001 Lights: Luciferins, Luciferases, Their Mechanisms of Action and Applications in Chemical Analysis, Biology and Medicine. Chem. Soc. Rev. 2016, 45 (21), 6048–6077. 10.1039/c6cs00296j

(42) Rautio, J.; Meanwell, N. A.; Di, L.; Hageman, M. J. The Expanding Role of Prodrugs in Contemporary Drug Design and Development. Nat. Rev. Drug Discov. 2018, 17 (8), 559–587. 10.1038/nrd.2018.46

(43) Zhang, R.; Qin, X.; Kong, F.; Chen, P.; Pan, G. Improving Cellular Uptake of Therapeutic Entities through Interaction with Components of Cell Membrane. Drug Deliv. 2019, 26 (1), 328–342. 10.1080/10717544.2019.1582730

(44) Rautio, J.; Kumpulainen, H.; Heimbach, T.; Oliyai, R.; Oh, D.; Järvinen, T.; Savolainen, J. Prodrugs: Design and Clinical Applications. Nat. Rev. Drug Discov. 2008, 7 (3), 255–270. 10.1038/nrd2468

(45) Lukowski, J. K.; Savas, C. P.; Gehring, A. M.; McKary, M. G.; Adkins, C. T.; Lavis, L. D.; Hoops, G. C.; Johnson, R. J. Distinct Substrate Selectivity of a Metabolic Hydrolase from Mycobacterium Tuberculosis. Biochemistry 2014, 53 (47), 7386– 7395. 10.1021/bi501108u

(46) Schué, M.; Maurin, D.; Dhouib, R.; N’Goma, J.-C. B.; Delorme, V.; Lambeau, G.; Carrière, F.; Canaan, S. Two Cutinase-like Proteins Secreted by Mycobacterium Tuberculosis Show Very Different Lipolytic Activities Reflecting Their Physiological Function. FASEB J. 2010, 24 (6), 1893–1903. 10.1096/fj.09-144766

(47) McKary, M. G.; Abendroth, J.; Edwards, T. E.; Johnson, R. J. Structural Basis for the Strict Substrate Selectivity of the Mycobacterial Hydrolase LipW. Biochemistry 2016, 55 (51), 7099–7111. 10.1021/acs.biochem.6b01057

(48) Ellis, E. E.; Adkins, C. T.; Galovska, N. M.; Lavis, L. D.; Johnson, R. J. Decoupled Roles for the Atypical, Bifurcated Binding Pocket of the ybfF Hydrolase. ChemBioChem 2013, 14 (9), 1134–1144. 10.1002/cbic.201300085

(49) Richter, L.; Tai, W.; Felton, J.; Saviola, B. Determination of the Minimal Acid-Inducible Promoter Region of the lipF Gene from Mycobacterium Tuberculosis. Gene 2007, 395 (1–2), 22–28. 10.1016/j.gene.2006.12.037

(50) Richter, L.; Saviola, B. The lipF Promoter of Mycobacterium Tuberculosis Is Upregulated Specifically by Acidic pH but Not by Other Stress Conditions. Microbiol. Res. 2009, 164 (2), 228–232. 10.1016/j.micres.2007.06.003

(51) Valente, E.; Simões, M. F.; Testa, B.; Constantino, L. Development of a Method to Investigate the Hydrolysis of Xenobiotic Esters by a *Mycobacterium Smegmatis* Homogenate. J. Microbiol. Methods 2011, 85 (2), 98–102. 10.1016/j.mimet.2011.02.003

(52) Ongwae, G. M.; Lepori, I.; Chordia, M. D.; Dalesandro, B. E.; Apostolos, A. J.; Siegrist, M. S.; Pires, M. M. Measurement of Small Molecule Accumulation into Diderm Bacteria. ACS Infect. Dis. 2023, 9 (1), 97–110. 10.1021/acsinfecdis.2c00435

(53) Fura, J. M.; Kearns, D.; Pires, M. M. D-Amino Acid Probes for Penicillin Binding Protein-Based Bacterial Surface Labeling *. J. Biol. Chem. 2015, 290 (51), 30540– 30550. 10.1074/jbc.M115.683342

(54) Liu, Z.; Lepori, I.; Chordia, M. D.; Dalesandro, B. E.; Guo, T.; Dong, J.; Siegrist, M. S.; Pires, M. M. A Metabolic-Tag-Based Method for Assessing the Permeation of Small Molecules Across the Mycomembrane in Live Mycobacteria. Angew. Chem. Int. Ed. 2023, 62 (20), e202217777. 10.1002/anie.202217777

(55) Pidgeon, S. E.; Apostolos, A. J.; Nelson, J. M.; Shaku, M.; Rimal, B.; Islam, M. N.; Crick, D. C.; Kim, S. J.; Pavelka, M. S.; Kana, B. D.; Pires, M. M. L,D-Transpeptidase Specific Probe Reveals Spatial Activity of Peptidoglycan Cross-Linking. ACS Chem. Biol. 2019, 14 (10), 2185–2196. 10.1021/acschembio.9b00427

(56) Johansen, M. D.; Herrmann, J.-L.; Kremer, L. Non-Tuberculous Mycobacteria and the Rise of Mycobacterium Abscessus. Nat. Rev. Microbiol. 2020, 18 (7), 392–407. 10.1038/s41579-020-0331-1

