## Supporting Information for "A Luciferase Reporter of Ester Prodrug Delivery to the Cytosol of Intracellular Mycobacteria"

### TABLE OF CONTENTS


| | |
|---|---|
| **SUPPLEMENTAL FIGURES** | S3-S9 |
| **Figure S1.** Calcein-AM treatment with E. coli in the presence of inhibitor of esterase activity. | S3 |
| **Figure S2.** Ethidium bromide (EtBr)-based membrane integrity assay of *Msm (wt)* compared to *Msm (luc)*. | S4 |
| **Figure S3.** Serum stability of D-luc ester derivatives incubated in mouse serum at 37 °C. | S5 |
| **Figure S4.** Luminescence of *Msm (luc)* cells upon addition of varying concentrations of D-luc-*n*Bu. | S6 |
| **Figure S5.** Ethidium bromide (EtBr)-based membrane integrity assay of *Msm (luc)* cells incubated with the D-luc ester series. | S7 |
| **Figure S6.** Luminescence of *E. coli (luc)* cells incubated with the D-luc ester series in the presence or absence of esterase co-expression. | S8 |
| **Figure S7.** Luminescence of *Msm (luc)* cells after the addition of D-luc was measured over 60 minutes in pH 7 vs pH 4. | S9 |
| **EXPERIMENTAL PROCEDURES** | S10-S16 |
| **Materials** | S10 |
| **Biological Methods** | S12 |
| Preparation of electrocompetent mycobacteria cells and transformation | S12 |
| Plasmid amplification and transformation of *E. coli* | S12 |
| Mycobacteria cell culture | S13 |
| *E. coli* cell culture | S13 |
| Mammalian cell culture | S13 |
| Bioluminescence-based accumulation assays | S13 |
| Bioluminescence-based accumulation assay of *Msm (luc)* in J774A.1 macrophages. | S14 |
| Cell-free luciferase assays | S14 |
| Cell-lysate assay | S15 |
| Ethidium bromide assay | S15 |
| Confocal microscopy analysis of PG-labeled Msm in J774A.1 macrophages. | S15 |
| Serum stability | S16 |
| Calcein-AM fluorescence assay | S16 |
| **Synthesis and Characterization of Compounds** | S17-S29 |
| General Methods | S17 |
| D-luc-Me | S19 |
| D-luc-Et | S21 |
| D-luc-*i*Pr | S23 |
| D-luc-*n*Bu | S26 |
| D-luc-*t*Bu | S29 |

### SUPPLEMENTAL FIGURES

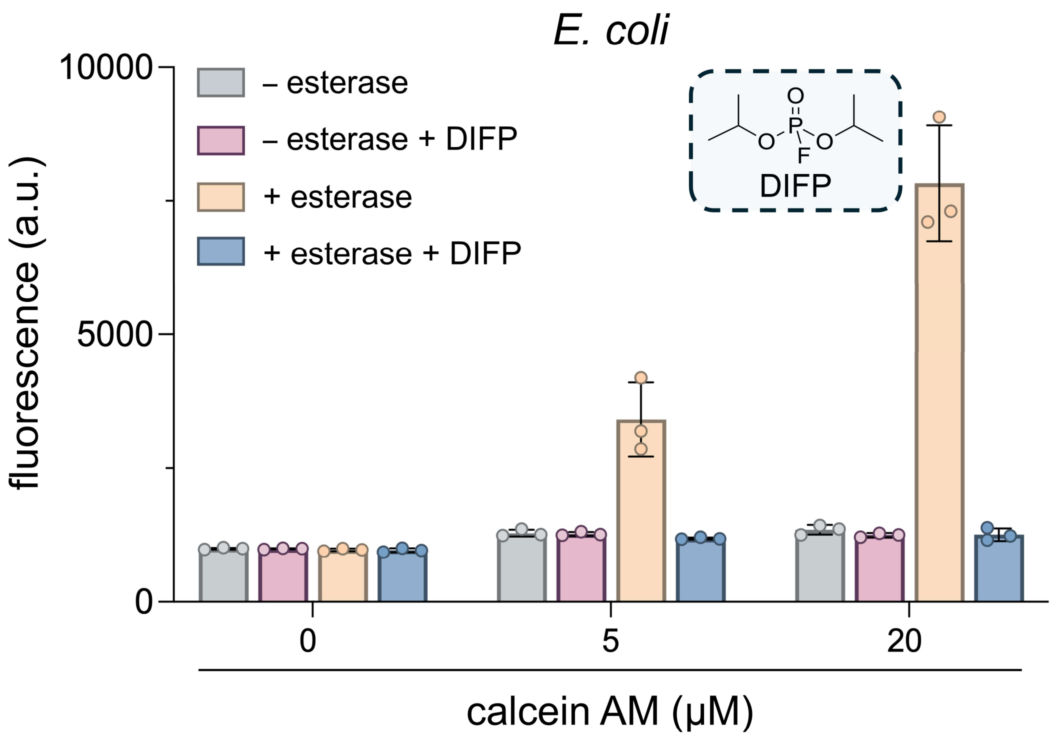


**Figure S1.** Esterase-dependent unmasking of calcein AM in *E. coli*. *E. coli* cells were treated with the indicated concentrations of calcein AM under four conditions: without expressing esterase, without expressing esterase and treated with DIFP, expressing esterase, and expressing esterase and treated with DIFP. DIFP (diisopropyl fluorophosphate, inset) irreversibly inhibits serine hydrolases. Fluorescence was measured after incubation and washing. Data are represented as mean ± SD ($n$ = 3).

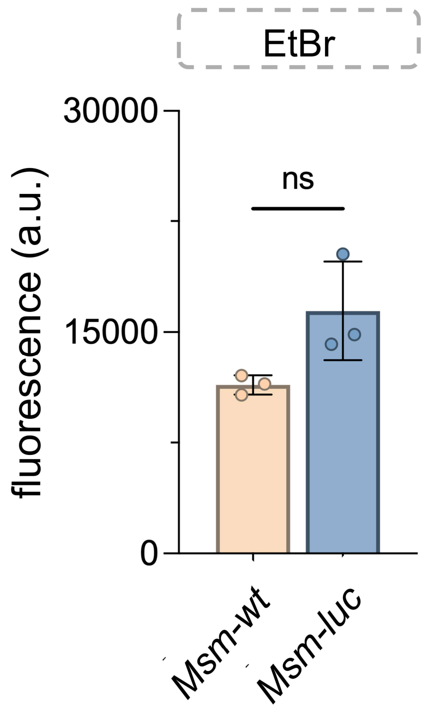


**Figure S2.** EtBr-based mycobacterial membrane integrity assay. *Msm (luc)* and *Msm (wt)* cells were incubated with EtBr (5 μM). Bars show fluorescence at the 60 min time point from the time-course data in Fig. 2e. Data are represented as mean ± SD (*n* = 3).

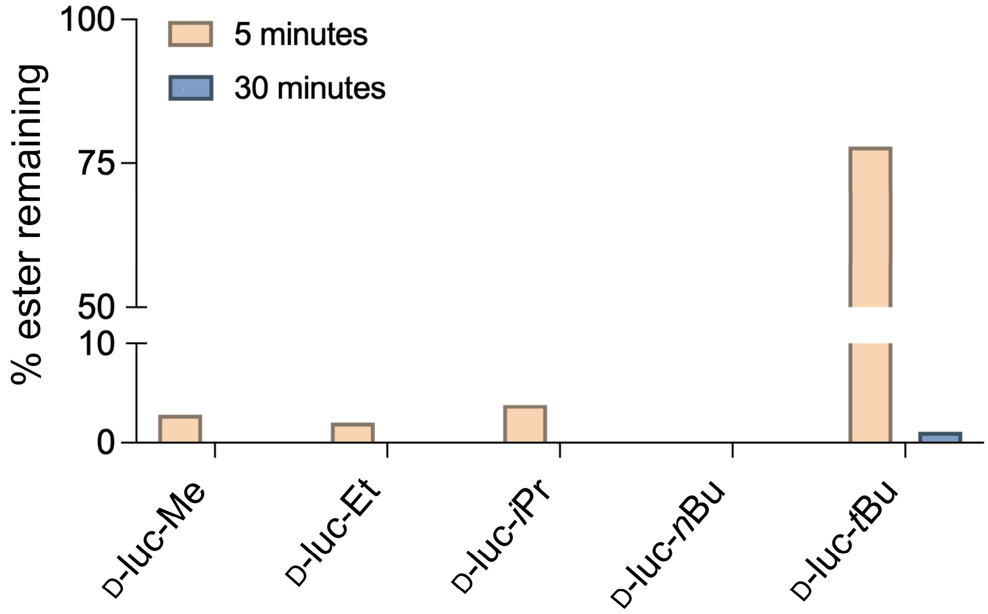


**Figure S3.** Comparison of serum stability of D-luc ester derivatives.

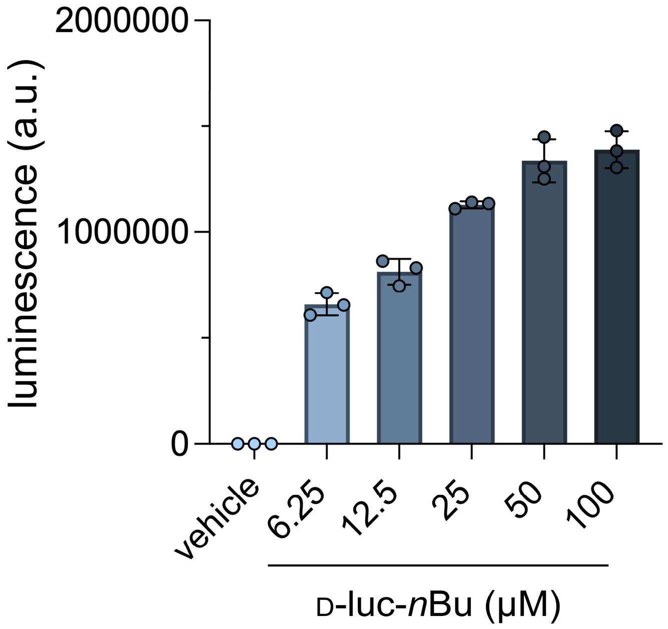


**Figure S4.** Luminescence of *Msm (luc)* cells immediately upon addition of varying concentrations of D-luc-*n*Bu. Data are represented as mean ± SD (*n* = 3).

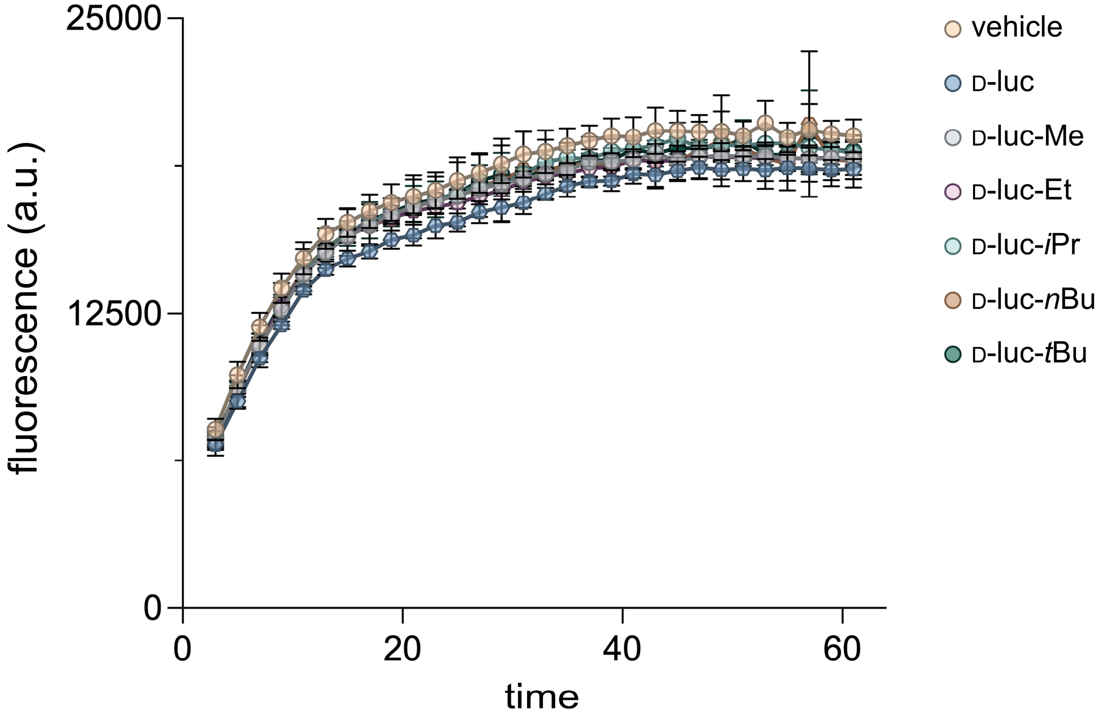


**Figure S5.** EtBr-based mycobacterial membrane integrity assay. *Msm (luc)* cells were co-incubated with EtBr (5 μM) and the D-luc ester derivatives. Fluorescence was monitored for over 60 minutes. Data are represented as mean ± SD (*n* = 3).

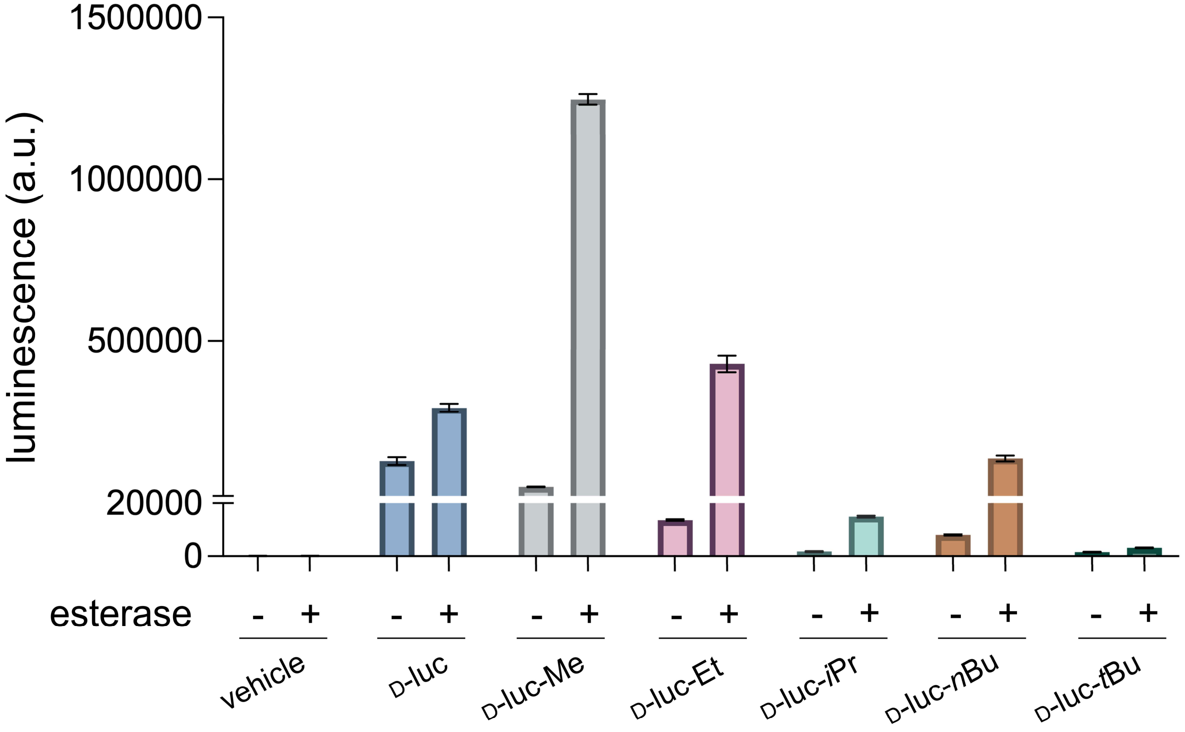


**Figure S6.** Luminescence of *E. coli (luc)* cells upon the addition of the D-luc ester series at the 60-minute time point in the presence or absence of co-expression of esterase. Data are represented as mean ± SD (*n* = 3).

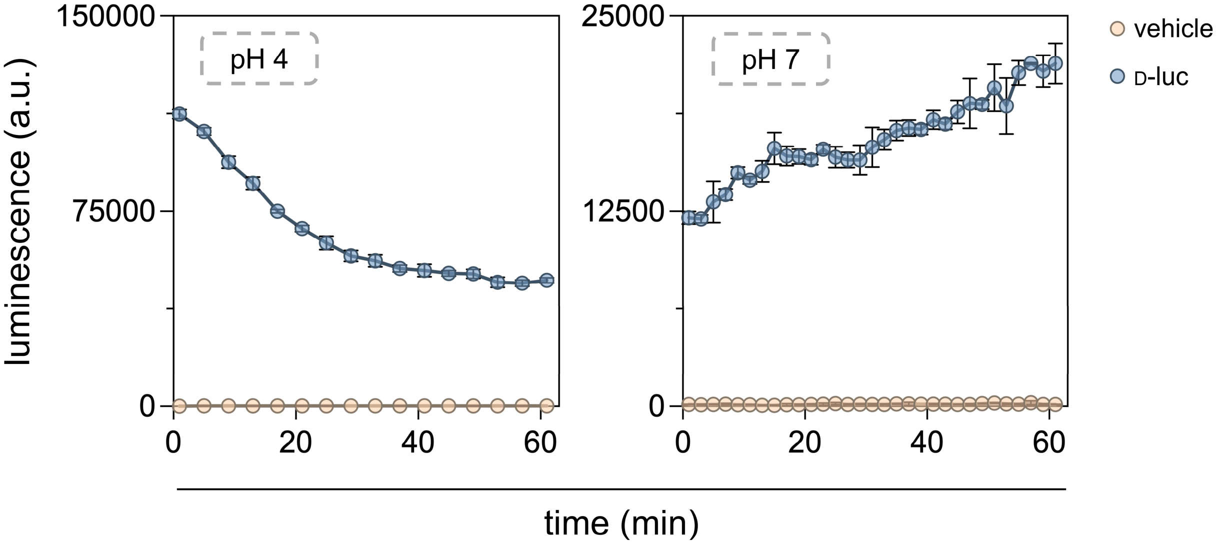


**Figure S7.** Luminescence of *Msm (luc)* cells after the addition of D-luc was measured over 60 minutes in pH 7 vs pH 4. Data are represented as mean ± SD ($n$ = 3).

### EXPERIMENTAL PROCEDURES

#### Materials

| REAGENTS | VENDOR SOURCE | CATALOG # |
|---|---|---|
| **Reagents for biological methods** | | |
| Middlebrook 7H9 media | VWR | 90003-876 |
| Catalase from bovine liver | Millipore Sigma | C1345 |
| D(+)-Glucose, anhydrous (Dextrose) | Chem Impex | 00805 |
| Bovine serum albumin fraction V | Millipore Sigma | 10735078001 |
| Glycerol | Millipore Sigma | G7893 |
| Tween 80 | VWR | 97061-674 |
| Tris Base | Fisher Scientific | BP152-1 |
| Ethidium bromide solution (10 mg/mL), Biotechnology Grade | VWR | 97064-970 |
| pMV306G13+FFluc | Addgene | 26157 |
| Electroporation Cuvettes, 0.2 cm gap | Bio-Rad | 652086 |
| QuantiLum® Recombinant Luciferase | Promega | E170A |
| Esterase from porcine liver | Millipore Sigma | E3019 |
| Dulbecco's Modified Eagle Medium (DMEM) | Fisher Scientific | 11885-084 |
| Fetal Bovine Serum | Fisher Scientific | A52568-01 |
| Penicillin-Streptomycin | Fisher Scientific | P4333 |
| T75 treated culture flasks | Fisher Scientific | 07-000-229 |
| Glass bottom 35 mm petri dishes | MatTek | P35GC-1.5-14-C |
| **Reagents for the synthesis and characterization of test molecules** | | |
| *N*, *N'*-Diisopropylethylamine (DIEA) | Chem Impex | 00141 |
| *N, N*-Dimethylformamide (DMF) ACS-grade, anhydrous | Millipore Sigma | 319937, 589565 |
| Dichloromethane (DCM) ACS-grade, anhydrous | Millipore Sigma | D65100, 270997 |
| Methanol ($CH_3OH$) ACS-grade, HPLC-grade | Millipore Sigma | 179337, 34860 |
| Acetonitrile ($CH_3CN$) ACS-grade, HPLC-grade | Millipore Sigma | 360457, 34851 |
| Anhydrous Tetrahydrofuran (THF) | Millipore Sigma | 186562 |
| Trifluoroacetic acid (TFA) HPLC grade | Millipore Sigma | 302031 |
| Trifluoroacetic acid (TFA) reagent-grade | Chem Impex | 00289 |
| **General materials and equipment** | | |

| | | |
|---|---|---|
| 96-well black flat bottom plates | Greiner | 655076 |
| 96-well clear untreated round bottom plates | VWR | 82050-622 |
| 4-Place Mini Bead Mill Homogeniser | VWR | 432-0366 |

### Biological Methods

**Preparation of electrocompetent cells and transformation of mycobacteria.**
*Mycobacterium smegmatis* mc²155 (*Msm*) and *Mycobacterium avium* clinical isolate 11 (MAH 11) were grown in 100 mL liquid medium to mid-log phase (OD600 = 0.5–0.7). Cells were harvested by centrifugation at 5000*g* for 10 min at 4°C in a Multifuge X Pro series centrifuge (Thermo Fisher Scientific) and resuspended in half the original culture volume of ice-cold 10% glycerol. The cells were washed several times with ice-cold 10% glycerol, concentrating two-fold with each wash, until a final volume of 2 mL was obtained. Competent cells were aliquoted into 200 µL portions and stored at −80°C.

For transformation, electroporation cuvettes were used with electroporation settings of 2.5 kV, 25 µF, and 200 Ω on a Bio-Rad gene pulser II. Cuvettes were pre-chilled on ice, and 100–150 µL of thawed competent cells were added per cuvette. One microgram of the firefly luciferase plasmid pMV306G13+FFluc (a gift from Brian Robertson & Siouxsie Wiles) was pipetted onto the cuvette wall before adding the cells to facilitate mixing. The cuvette exterior was dried, placed in the electroporator, and pulsed under the preset conditions.

Immediately after electroporation, cells were recovered by transferring them as quickly as possible into a vial containing pre-warmed, antibiotic-free 7H9 medium. Cultures were incubated for 3 h (for *Msm*) or overnight (for *Mav*) at 37°C with shaking, then pelleted by centrifugation. The cell pellet was resuspended in 7H9 and 100 µL of the suspension was plated as is, while the rest of the culture was plated after centrifugation to concentrate cells. Culturing of the resulting colonies on 50 µg/mL kanamycin-containing agar plates yielded *Msm (luc)* and *Mav (luc).*

**Plasmid amplification and transformation of *E. coli*.**
The FLUC2 pET28a plasmid was obtained from the Ai laboratory at the University of Virginia, and the pET21a(+) plasmid encoding *Pseudomonas fluorescens* carboxylesterase 2 was obtained from GenScript. For plasmid amplification, 50 µL of competent DH5α *E. coli* cells and 2–5 µL of FLUC2 pET28a plasmid DNA were mixed and incubated on ice for 30 min. Cells were heat-shocked at 42 °C for 30 s, followed by another 2 min on ice. One mL of sterile LB medium was added, and cells were incubated at 37 °C for 1 h. Subsequently, 25–200 µL of the recovered cells were plated on LB agar containing kanamycin (50 µg/mL) and incubated at 37 °C overnight. Individual colonies were picked and grown overnight in LB broth containing kanamycin (50 µg/mL). The following day, plasmid DNA was isolated using a ZymoPURE Plasmid Miniprep Kit (Zymo Research). The same procedure was used to amplify and isolate the carboxylesterase 2 pET21a(+) plasmid, with ampicillin (50 µg/mL) replacing kanamycin during selection and overnight culture.

For transformation, the isolated FLUC2 pET28a plasmid was introduced into chemically competent BL21(DE3) *E. coli* cells using the heat-shock and recovery conditions described above. Culturing of the transformed cells on kanamycin-containing LB agar plates (50 µg/mL) yielded *E. coli (luc)*. For simultaneous co-expression, the FLUC2 pET28a and carboxylesterase 2 pET21a(+) plasmids were added to the same aliquot of chemically competent BL21(DE3) *E. coli* cells, followed by heat shock and recovery as described above. Culturing of the transformed cells on LB agar plates containing kanamycin (50 µg/mL) and ampicillin (50 µg/mL) yielded *E. coli (luc + est).*

**Mycobacteria cell culture.**

*Msm (luc)* cells were grown in 7H9 media with 0.5% glycerol, 0.05% Tween 80, and 1x ADC enrichment. 10x ADC was made with 5 g bovine serum albumin, 2 g dextrose, 3 mg catalase in 100 mL autoclaved milliQ $H_2O$. Glycerol stocks were prepared from stationary-phase cells in 30% glycerol, and aliquots were stored at −80°C. Unless otherwise mentioned, cells were inoculated from glycerol stocks into 7H9 medium at a 1:100 dilution in standard culture tubes and grown for 22 h at 37°C until reaching an $OD_{600}$ of approximately 1.0. Whenever appropriate, *Msm (wt)* cells were also grown to $OD_{600}$ of approximately 1.0 as a control. Cells were then harvested by centrifugation at 3000 × g for 2 min in a Heraeus Multifuge X1 centrifuge (Thermo Fisher Scientific), washed twice with phosphate-buffered saline containing 0.05% Tween 80 (PBST, pH 7.4) and resuspended in PBST for subsequent assays.

***E. coli* cell culture.**

*E. coli (luc)* cells were grown in LB medium containing kanamycin (50 µg/mL), while *E. coli (luc + est)* cells were grown in LB medium containing kanamycin (50 µg/mL) and ampicillin (50 µg/mL). Glycerol stocks were prepared by mixing equal volumes of overnight culture and 60% glycerol in water, yielding a final glycerol concentration of 30%. Unless otherwise mentioned, cells were inoculated from glycerol stocks into 3 mL of LB medium containing the corresponding antibiotics in standard culture tubes and grown overnight at 37 °C with shaking. Cultures were then diluted 1:10 into fresh LB medium containing the same antibiotics and grown at 37 °C until reaching an $OD_{600}$ of 0.6–0.8. Protein expression was induced by adding isopropyl β-D-1-thiogalactopyranoside (IPTG) to a final concentration of 1 mM, followed by incubation at 37 °C for 2 h with shaking. Cells were then harvested by centrifugation at 3000 × g for 3 min, washed twice with phosphate-buffered saline (PBS, pH 7.4), and resuspended in PBS for subsequent assays.

**Mammalian cell culture.**

J774A.1 cells were cultured in Dulbecco's modified Eagle's medium (DMEM) supplemented with 10% (v/v) FBS, 50 IU/mL penicillin, and 50 µg/mL streptomycin in a T75 flask and maintained in a humidified atmosphere of 5% $CO_2$ at 37°C.

**Bioluminescence-based accumulation assays.**

In a 96-well, black, flat-bottomed plate, molecules of interest were added to achieve the desired final concentrations in a total volume of 100 µL. Unless otherwise mentioned, D-luc and its ester derivatives were assessed at a concentration of 25 µM. Stock solutions of these molecules were prepared in PBS. PBS was also added to blank and negative control wells to maintain equal total volumes and cell numbers across all wells.

Equal volumes of cells washed and resuspended in PBS were then transferred to each well and mixed. For assays designed to mimic phagosomal pH, cells were resuspended in a phosphate-citrate buffer consisting of 0.2 M disodium phosphate and 0.1 M citric acid, with the pH adjusted to 4.5. The plate was then placed in an Agilent BioTek Synergy H1 Microplate Reader (or BioTek Synergy2 plate reader for *Mav*) set to kinetic luminescence mode, typically recording every 1 min over 60 min. During all luminescence measurements, continuous slow double-orbital shaking at 237 cpm was applied. Luminescence was measured using fiber optics, with the gain set to 100, integration time of 0:01:00, and a 100 ms delay after plate movement. The read height was 4.5 mm, and the temperature was maintained at 37 °C. For experiments involving multiple strains, $OD_{600}$ was normalized between treatment arms to ensure consistent cell numbers across all conditions.

**Bioluminescence-based accumulation assay of *Msm (luc)* in J774A.1 macrophages.**

*Msm (luc)* cells were grown in liquid medium as described above for approximately 24 hours to mid-log phase ($OD_{600}$ = 0.5-0.7) while shaking (250 rpm) at 37°C. The bacteria were harvested and centrifuged for 2 min at 4000 x g then washed three times with PBST. J774A.1 cells were cultured as described above. On the day prior to the experiment, J774A.1 cells were seeded into a T-75 flask and allowed to adhere. On the day of the experiment, J774A.1 cells were washed with twice with 1X PBS. *Msm (luc)* cells were resuspended in DMEM + 10% FBS containing no antibiotics and added to the washed J774A.1 cells at an MOI of 100. The cell mixture was then incubated at 37°C for 1 hour to induce phagocytosis, then washed five times with 1X PBS to remove extracellular bacteria. Cells were then removed from the flask by scraping, centrifuged for 5 min at 1100 x g, and resuspended in appropriate volume of 1X PBS such that aliquoting 95 µL of the suspension to each well required in the plate would yield approximately $4x10^5$ cells per well. A 96-well, black, flat-bottomed plate was prepared with 5 µL of the molecules of interest as described above. 95 µL of the prepared cell suspension was added to each well, mixed, and the plate was placed in an Agilent BioTek Synergy H1 Microplate Reader set to kinetic luminescence mode, recording every 1 min over 60 min. During luminescence measurements, continuous slow double-orbital shaking at 237 cpm was applied. Luminescence was measured using fiber optics, with the gain set to 200, integration time of 0:01:00, and a 100 ms delay after plate movement. The read height was 4.5 mm, and the temperature was maintained at 37 °C.

**Cell-free luciferase assays.**

The 14.9 mg/mL luciferase enzyme stock (QuantiLum® Recombinant Luciferase received from Promega) was diluted in 25 mM Tris buffer (pH 8) to prepare 0.4 mg/mL aliquots (40 µL each), which were stored at −80 °C. All assays were conducted in 25 mM Tris buffer (pH 8) in 96-well, black, flat-bottomed plates. Molecules of interest were added to achieve the desired final concentrations in a total volume of 100 µL. Appropriate volumes of $MgSO_4$ (100 mM), freshly prepared ATP (20 mM), and luciferase stock (0.4 mg/mL) were added to yield final concentrations of 5 mM, 1 mM, and 20 µg/mL, respectively. Where indicated, porcine liver esterase (PLE) was added to yield a final concentration of 1.5 U/mL. Plates were placed in an Agilent BioTek Synergy H1 Microplate Reader set to kinetic luminescence mode, typically recording every 2 min for 60 min. Continuous slow double-orbital shaking at 237 cpm was applied during all measurements. Luminescence was recorded using fiber optics, with gain set to 100, integration time of 0:01:00, a 100 ms delay after plate movement, read height of 4.5 mm, and temperature maintained at 37 °C.

For cell-free assays with esterase, the 14.9 mg/mL luciferase enzyme stock (QuantiLum® Recombinant Luciferase received from Promega) was diluted in 25 mM Tris buffer (pH 8) to prepare 0.4 mg/mL aliquots (40 µL each), which were stored at −80 °C. Stocks of porcine liver esterase (Sigma) were prepared in the same buffer at 5 U/µL. All assays were conducted in 25 mM Tris buffer (pH 8) in 96-well, black, flat-bottomed plates. Compounds were added to achieve the desired final concentrations in a total volume of 100 µL. All treatments contained $MgSO_4$ (5 mM), freshly prepared ATP (1 mM), and luciferase stock (20 µg/mL) and were done in triplicate. For select treatments, esterase was added to a final concentration of 0.2 U/µL. Plates were placed in an Agilent BioTek Synergy H1 Microplate Reader at 37 °C with continuous slow double-orbital shaking at 237 cpm. Luminescence was recorded every 1 min for 60 min with gain set to 100, integration time of 0:01:00, a 100 ms delay after plate movement, and read height of 4.5 mm.

**Cell-lysate assay.**
This procedure was adapted from Tallman et al., *ACS Infect. Dis.,* 2016. Cell pellets were thawed and resuspended in lysis buffer consisting of 50 mM Tris (pH 7.5 at 4 °C), 200 mM NaCl, 0.5 mM $CaCl_2$, 0.5 mM $MgCl_2$, and 0.2% Triton X-100. Cells were lysed by mechanical disruption (VWR®, 4-Place Mini Bead Mill Homogeniser) using zirconia/silica beads in screw-cap tubes (three 1-min pulses with intermittent cooling on ice). Cell debris and beads were removed by centrifugation at 13,000 × g for 10 min at 4 °C, and the clarified supernatant was transferred to a fresh tube. The lysate was further clarified by centrifugation at 15,700 × g for 5 min at 4 °C to remove residual insoluble material. Final lysates were sterilized by filtration through a 0.2 µm PVDF membrane.

**Ethidium bromide assay.**
In a 96-well, black, flat-bottomed plate, molecules of interest were added in triplicate to achieve the desired final concentrations in a total volume of 100 µL. To each well, 5 µL of 100 µM ethidium bromide (EtBr) was added to achieve a final concentration of 5 µM. *Msm (luc)* cells were then added to each well to make up the volume. The plate was placed in an Agilent BioTek Synergy H1 Microplate Reader set to kinetic fluorescence mode, recording every 2 min over 60 min. Fluorescence was measured with excitation and emission wavelengths of 530 nm and 590 nm, respectively. Continuous slow double-orbital shaking at 237 cpm was applied, and the temperature was maintained at 37 °C.

**Confocal microscopy analysis of PG-labeled *Msm* in J774A.1 macrophages.**
*Msm (wt)* cells were grown in liquid medium as described above for approximately 24 hours to mid-log phase ($OD_{600}$ = 0.5-0.7) while shaking (250 rpm) at 37°C. At this time, 100 µM of DADA was added to the media and the cells were further grown for 16 h at 37°C while shaking (250 rpm) to a stationary phase. The bacteria were harvested and

centrifuged for 2 min at 4000 x g then washed three times with PBST. J774A.1 cells were cultured as described above. On the day prior to the experiment, J774A.1 cells were seeded into 35 mm glass bottom microwell dishes and allowed to adhere. On the day of the experiment, J774A.1 cells were washed twice with 1X PBS. The PG-labeled *Msm* cells were resuspended in DMEM + 10% FBS containing no antibiotics and added to the washed J774A.1 cells at an MOI of 100. The cell mixture was then incubated at 37°C for 1 hour to induce phagocytosis. The cell mixture was washed five times with 1X PBS to remove extracellular bacteria and fixed for 30 min with 4% formaldehyde in 1X PBS. Cells were imaged using a Leica STELLARIS 8 Super resolution microscopy system (40x/1.3 oil-immersion lens) equipped with 405 nm laser, emission gated from 420 – 480 nm. We acknowledge the Keck Center for Cellular Imaging for the usage of the Leica STELLARIS 8 microscopy system. [PI: AP; NIH-OD030409].

**Serum stability.**
Compounds were incubated in mouse serum at a concentration of 0.5 mM to a total of 200 µL at 37 °C. At each time point (0, 15, 30, 60 min), 40 µL of the solution was aliquoted and diluted in a 1:1 solution of $H_2O/CH_3CN$ (101 µL) containing trichloroacetic acid (9 µL) and 100 µM of internal standard Fmoc-L-phenylalanine. This mixture was incubated in ice for 5 min to precipitate serum proteins. The samples were then centrifuged at 21,000×g for 5 min. 140 µL of the supernatant was then transferred to a filter tube and centrifuged again at 21,000×g for 5 min to filter. 110 µL of the filtrate was then analyzed using Phenomenex Luna 5 µm C8(2) on RP-HPLC; gradient elution in $H_2O/CH_3CN$ with 0.1% TFA at 1 mL/min. Absorbance was monitored at 330 nm (D-luciferin) and 254 nm (Fmoc-L-phenylalanine). The area under the curve (AUC) for the peaks corresponding to Fmoc-L-phenylalanine, the luciferin esters, and any luciferin generated from ester hydrolysis was obtained using Waters® Empower software. The AUCs of luciferin and the luciferin esters were normalized to the AUC of Fmoc-L-phenylalanine from each run. The data was plotted by dividing each ester signal by the combined ester + luciferin signal from each run.

**Calcein-AM fluorescence assay.**
*E. coli* and *Msm* cells were cultured, harvested, and washed as described above. Cells were resuspended in PBS and normalized to equal $OD_{600}$ values. Calcein acetoxymethyl ester (calcein-AM) was added to the cell suspensions at the indicated final concentrations. Where indicated, cells were co-treated with calcein-AM and the serine hydrolase inhibitor diisopropyl fluorophosphate (DIFP) at a final concentration of 100 µM. Following incubation for 1 h at 37 °C, cells were washed three times with PBS and resuspended in PBS. Cell suspensions were transferred to 96-well, black, flat-bottomed plates, and fluorescence was measured using an Agilent BioTek Synergy H1 Microplate Reader with excitation and emission wavelengths of 494 nm and 517 nm, respectively.

### Synthesis and Characterization of Compounds

#### General Methods

For the synthesis of D-luc-Me, D-luc-Et, D-luc-*i*Pr, and D-luc-*n*Bu, D-luciferin (50 mg, 0.179 mmol) was dissolved in the appropriate alcohol containing the corresponding desired ester group (6.0 mL, excess) and stirred for 1 h at room temperature, achieving partial dissolution. Thionyl chloride (80 µL, 1.11 mmol, 6.2 eq.) was then carefully added dropwise *via* a syringe over a few minutes, and the resulting reaction mixture was stirred overnight, becoming clear yellowish orange in color. Volatiles were subsequently removed under reduced pressure, and the leftover residue was partitioned between EtOAc and $H_2O$. The aqueous phase was adjusted to pH 6, and the organic phase was separated, washed with $H_2O$ (2x) and brine, and dried over $Na_2SO_4$, filtered, and concentrated under reduced pressure to afford the corresponding ester. D-luc-*i*Pr, and D-luc-*n*Bu were further purified by reverse-phase preparative high-performance liquid chromatography (RP-HPLC) equipped with Waters 1525 with 2489 UV/Visible Detector on a Phenomenex Luna 10 µm C8(2) 100 Å (250 x 21.2 mm) column using gradient elution with $H_2O$/MeOH with 0.1% TFA at 10 mL/min. The HPLC fractions of the desired purified compounds were first concentrated under reduced pressure using a rotary evaporator. The final concentrated aqueous solutions were lyophilized to dryness using Labconco Freezone 4.5 L (-84 $^oC$) lyophilizer.

For the synthesis of D-luc-*t*Bu, D-Luciferin (40 mg, 0.143 mmol) was dissolved in tert-butyl acetate (5.0 mL) and cooled to 0 °C. Bistriflimide (44 mg, 0.157 mmol, 1.1 equiv) was then added, and the reaction mixture was stirred for 4h at 0 °C. The reaction was quenched with saturated $NaHCO_3$, and the mixture was extracted with $CH_2Cl_2$ (3 X, 10 mL). The combined organic layers were dried over $Na_2SO_4$, filtered, and concentrated under reduced pressure to afford a yellow residue. The crude product was purified by reverse-phase preparative high-performance liquid chromatography (RP-HPLC) equipped with Waters 1525 with 2489 UV/Visible Detector on a Phenomenex Luna 10 µm C8(2) 100 Å (250 x 21.2 mm) column using gradient elution with $H_2O$/MeOH with 0.1% TFA at 10 mL/min. The HPLC fractions of the desired purified compounds were first concentrated under reduced pressure using a rotary evaporator. The final concentrated aqueous solutions were lyophilized to dryness using Labconco Freezone 4.5 L (-84 $^oC$) lyophilizer to afford D-luc-*t*Bu as a white solid (7.9 mg, 15%).

Compounds were analyzed for purity on a Waters 1525 with 2489 UV/Visible Detector RP-HPLC using Phenomenex Luna 5 µm C8(2); gradient elution in $H_2O/CH_3OH$ with 0.1% TFA at 1 mL/min. Analytical HPLC indicated >95% purity for all compounds. Electrospray ionization (ESI) mass spectrometry was performed on an Advion Expression® CMS using standard parameters. A low-fragmentation, low-energy setup was employed for fragmentation-sensitive samples. HRMS analysis was performed on Agilent 6545B Q-TOF mass spectrometer equipped with LC 1260 Infinity II LC system with auto sampler and C18(2) column (Luna 5 µm 100Å 250 x 4.6 mm. Mass m/z detection was observed using purine and hexakis(1H,1H,3H-tetrafluoropropoxy)phosphazine as internal standard. 1H and 13C NMR spectra of final compounds and intermediates were obtained on a Varian 600 MHz spectrometer. All spectra were processed and analyzed using MestreNova software. Residual solvent signals from deuterated solvents were used as an internal standard with reference to tetramethylsilane (TMS) for defining chemical shifts. Chemical shifts are reported in parts per million or ppm ($\delta$) and coupling constants ($J$) are reported in hertz (Hz).

**D-luc-Me**

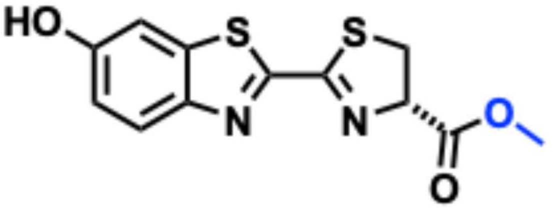

**Molecular formula:** $C_{12}H_{10}N_2O_3S_2$. **HRMS:** *m/z* calcd for $[M+H]^+$, 295.0206; found, 295.0206. $^1$**H NMR** (DMSO-d6) δ 10.2 (s, 1H, OH), 7.96 (d, 1H, ArH), 7.44 (d, 1H, ArH), 7.06 (dd, 1H, ArH), 5.51 (1H, N–CH), 3.79 9dd, 1H, S-$CH_2$), 3.75 (s, 3H, $OCH_3$), 3.67 (dd, 1H, S–$CH_2$). NMR has a trace amount of grease impurity (~1.2ppm).

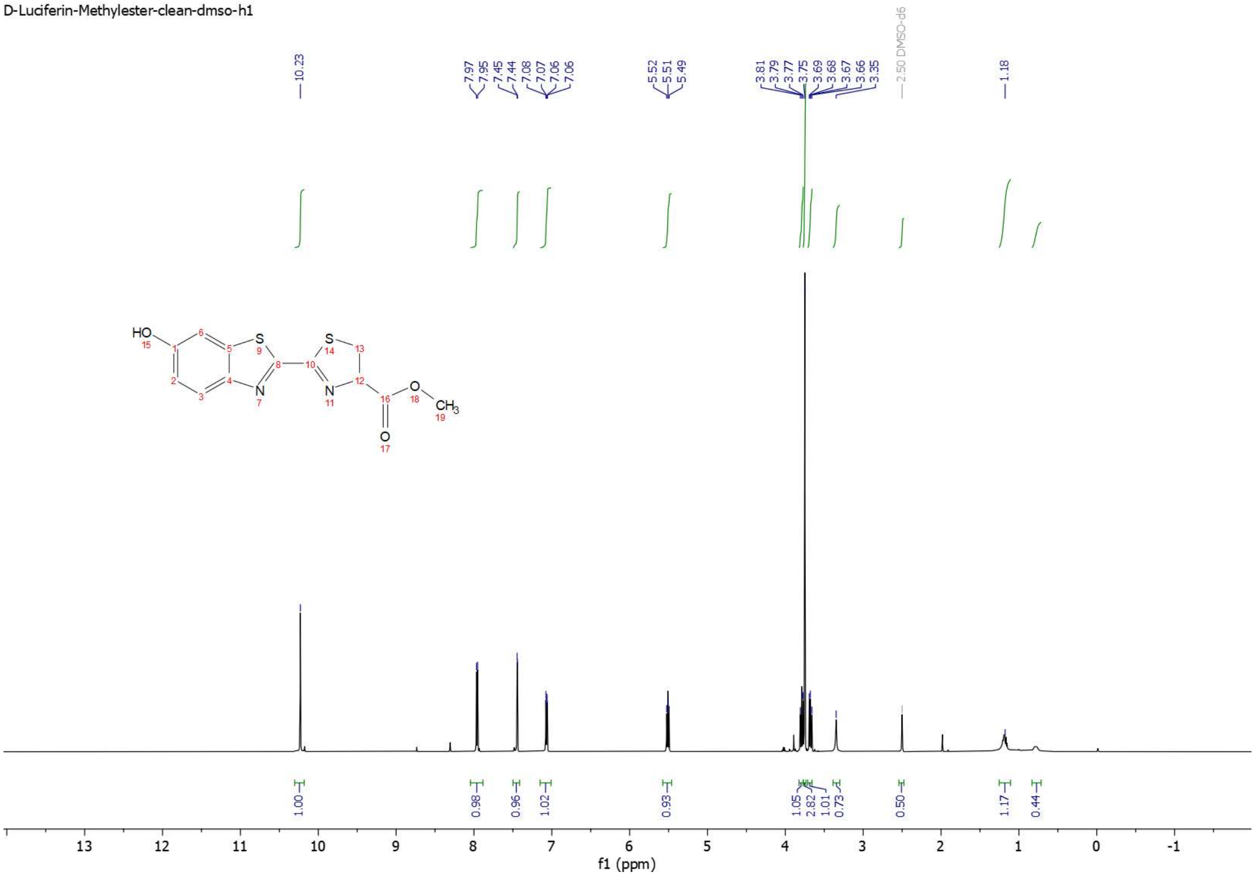


$^1$H NMR spectrum of D-luc-Me.

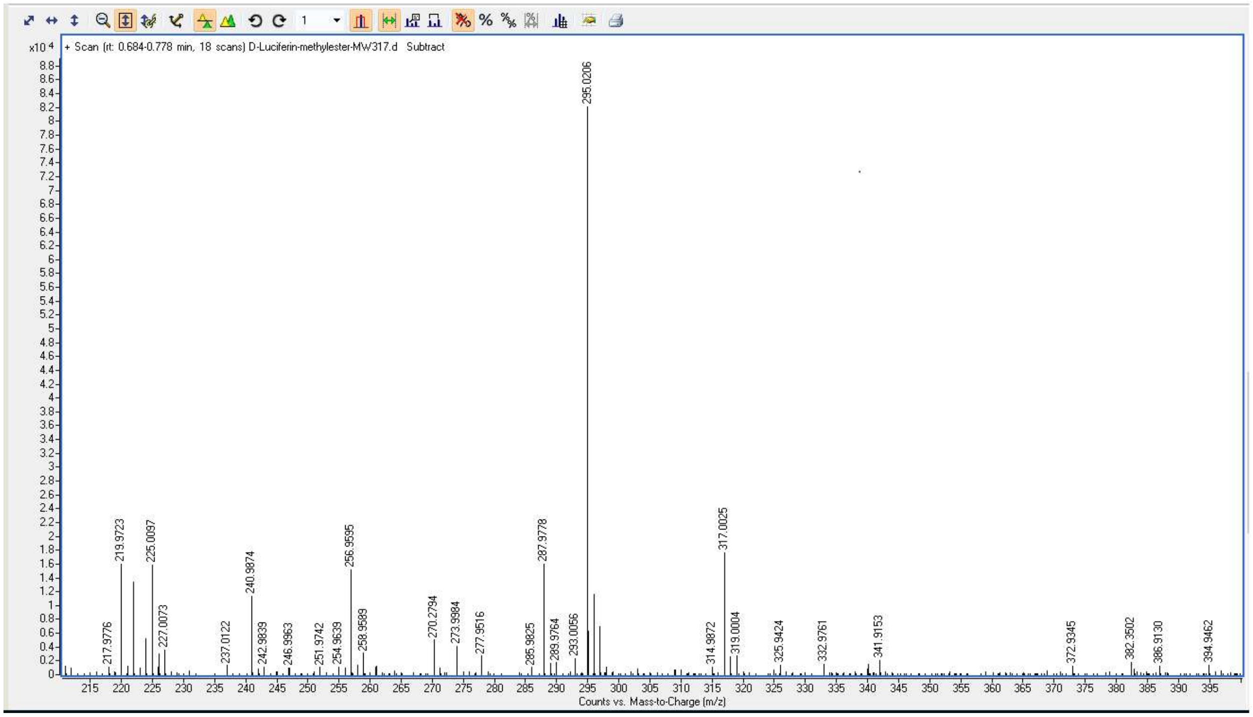


High Resolution Mass Spectrum for D-luc-Me (expected m/z 295.0206 for $[M+H]^+$).

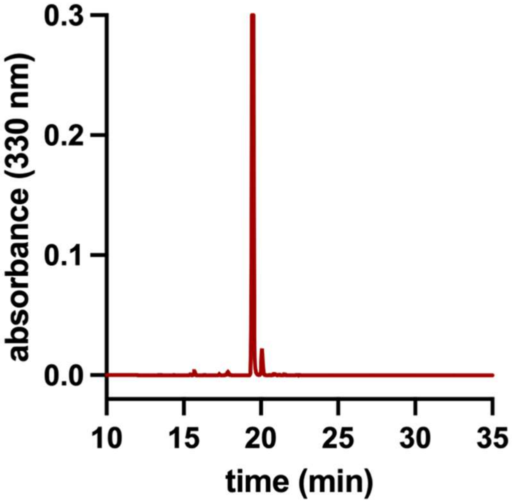


Analytical HPLC Chromatogram of D-luc-Me.

**D-luc-Et**

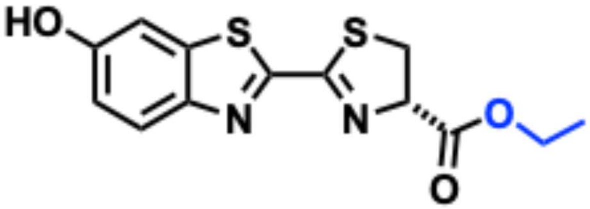


**Molecular formula:** $C_{13}H_{12}N_2O_3S_2$. **HRMS:** *m/z* calcd for $[M+H]^+$, 309.0362; found, 309.0365. **$^1$H NMR** (DMSO-$d_6$) δ 10.22 (s, 1H, OH), 7.95 (d, 1H, ArH), 7.44 (s, 1H, ArH), 7.06 (d, 1H, ArH), 5.48 (t, 1H, N–CH), 4.20 (t, 2H, O–$CH_2$), 3.79 (dd, 1H, S–$CH_2$), 3.66 (dd, 1H, S–$CH_2$), 1.25 (t, 3H, $CH_3$).

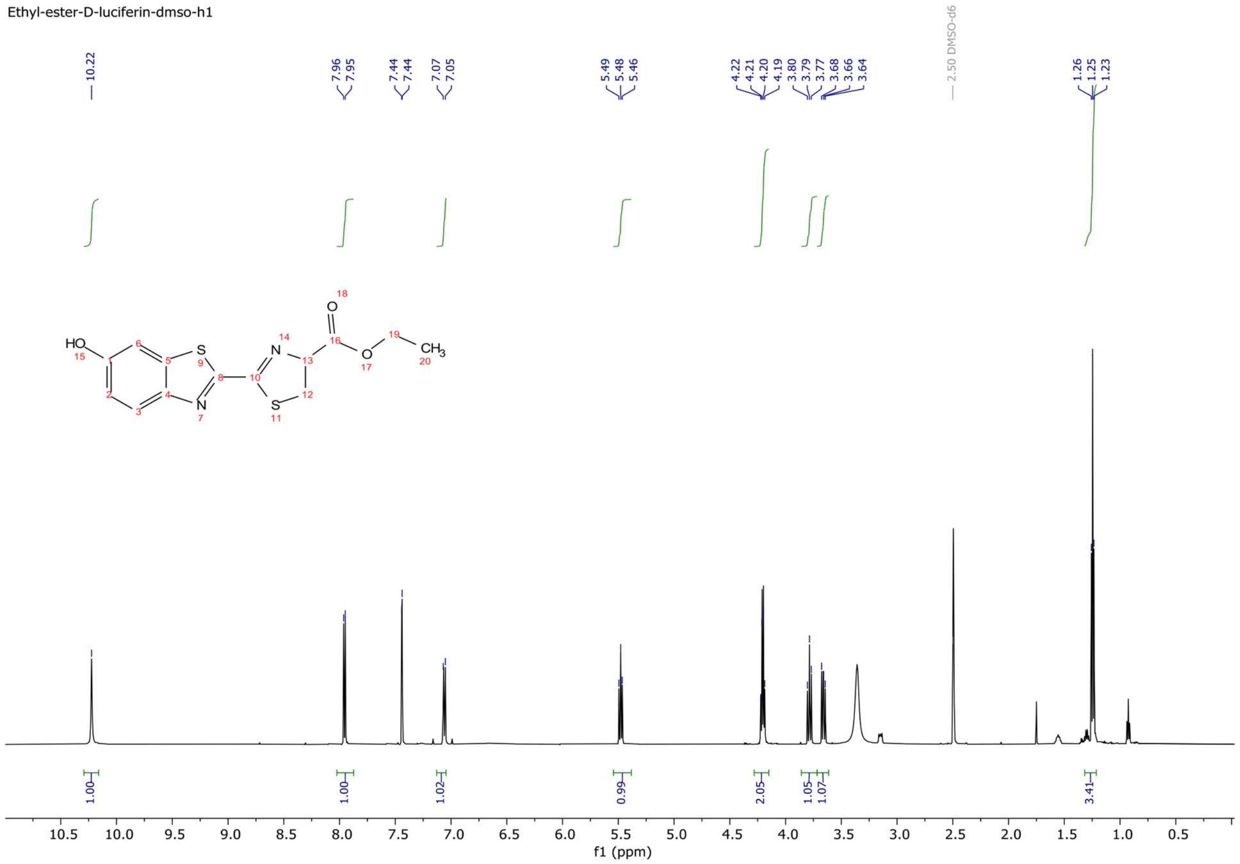


$^1$H NMR spectrum of D-luc-Et.

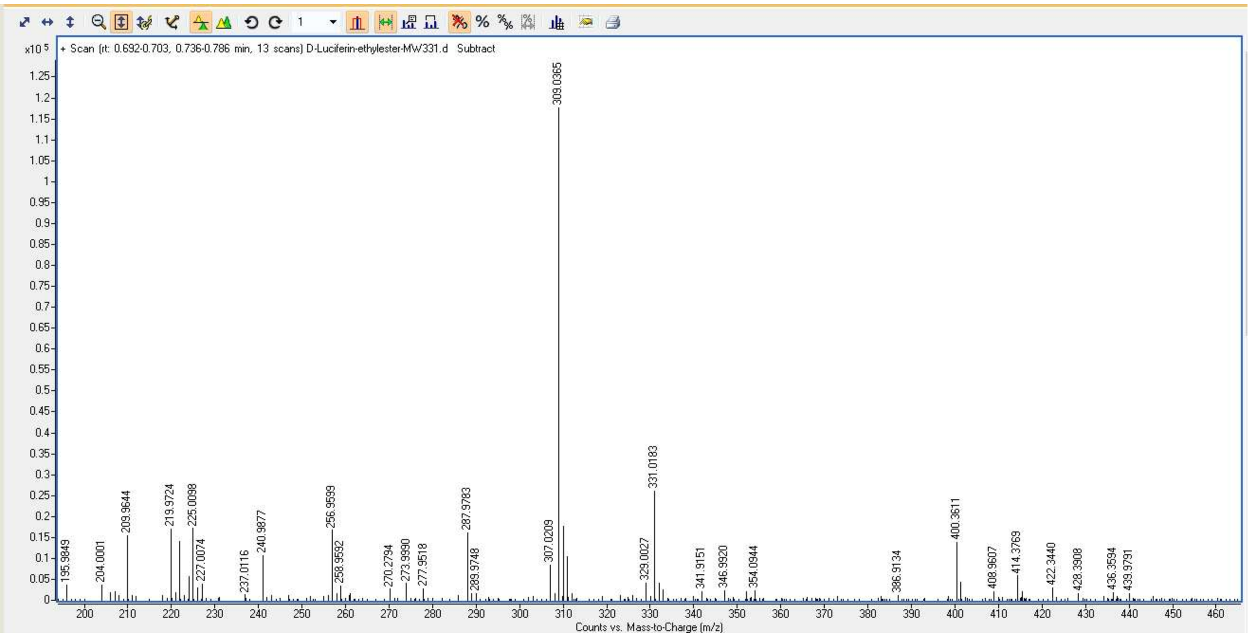

High Resolution Mass Spectrum for D-luc-Et (expected m/z 309.0362 for $[M+H]^+$).

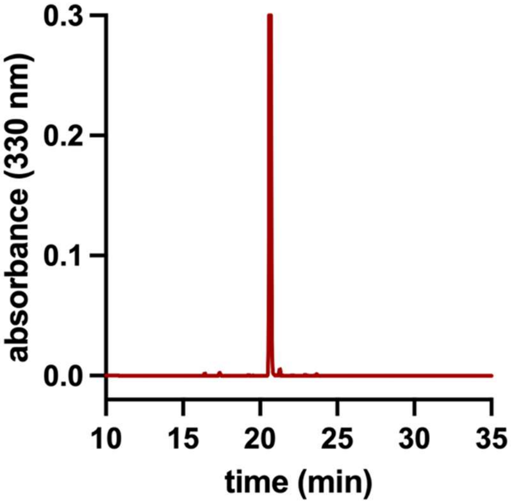

Analytical HPLC Chromatogram of D-luc-Et.

**D-luc-*i*Pr**

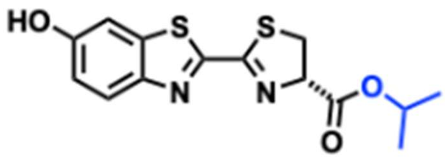

**Molecular formula:** $C_{14}H_{14}N_2O_3S_2$. **HRMS:** *m/z* calcd for $[M+H]^+$, 323.0519; found, 323.0523. **$^1$H NMR** (DMSO-$d_6$) δ 10.23 (s, 1H, OH), 7.96 (d, 1H, ArH), 7.44 (s, 1H, ArH), 7.07 (d, 1H, ArH), 5.44 (t, 1H, N–CH), 5.00 (m, 1H, O–CH), 3.78 (dd, 1H, S–$CH_2$), 3.64 (dd, 1H, S–$CH_2$), 1.25 (d, 6H, $(CH_3)_2$). **$^{13}$C NMR** (DMSO-$d_6$) δ 169.4, 165.0, 157.4, 156.6, 146.3, 137.2, 124.9, 117.2, 106.8, 77.9, 69.0, 34.7, 21.5.

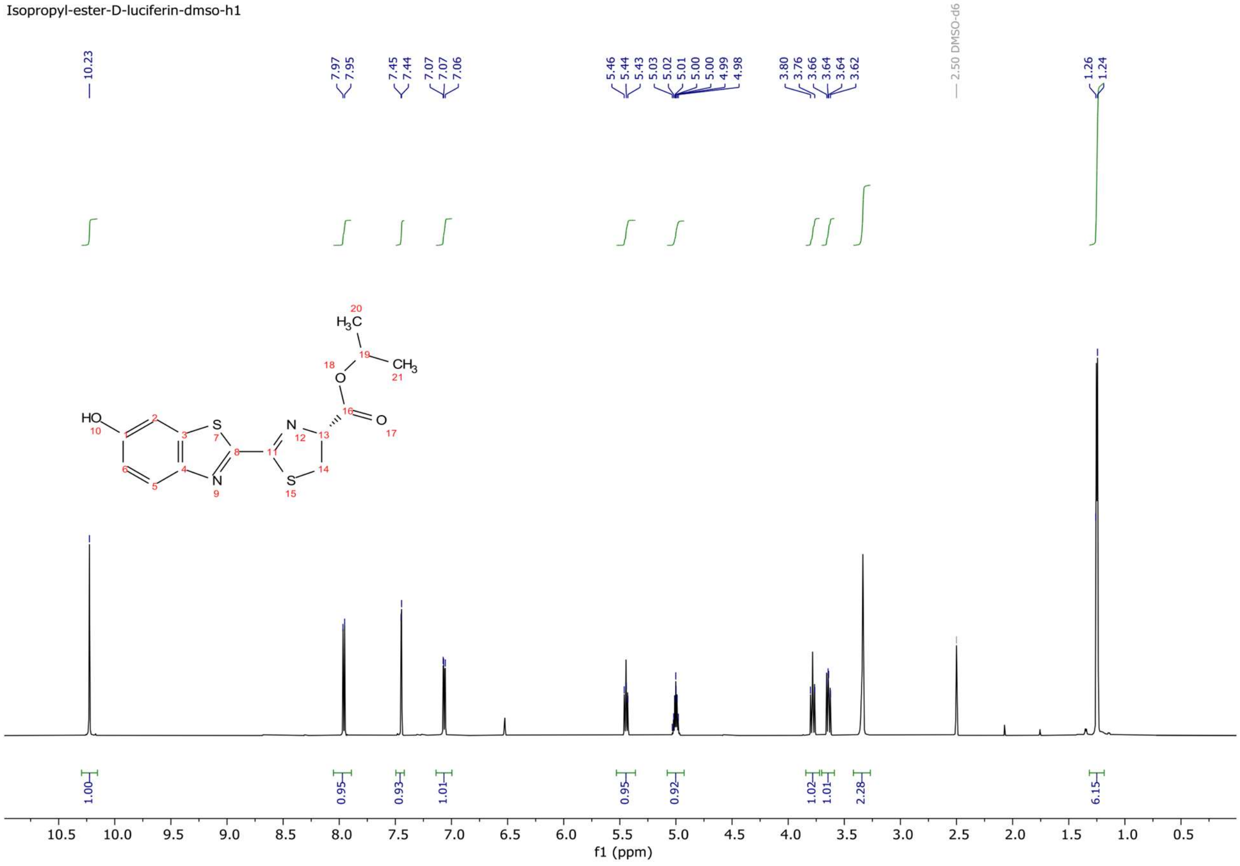


$^1$H NMR spectrum of D-luc-*i*Pr.

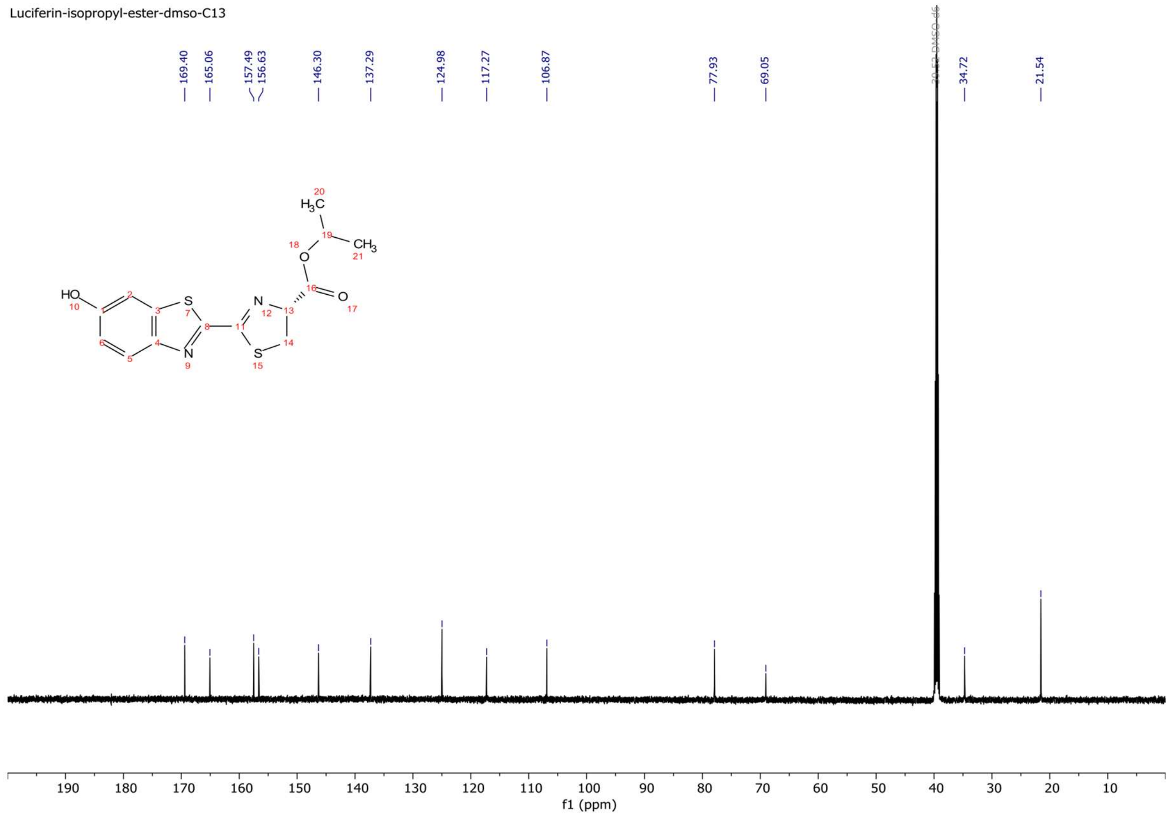


$^{13}$C NMR spectrum of D-luc-*i*Pr.

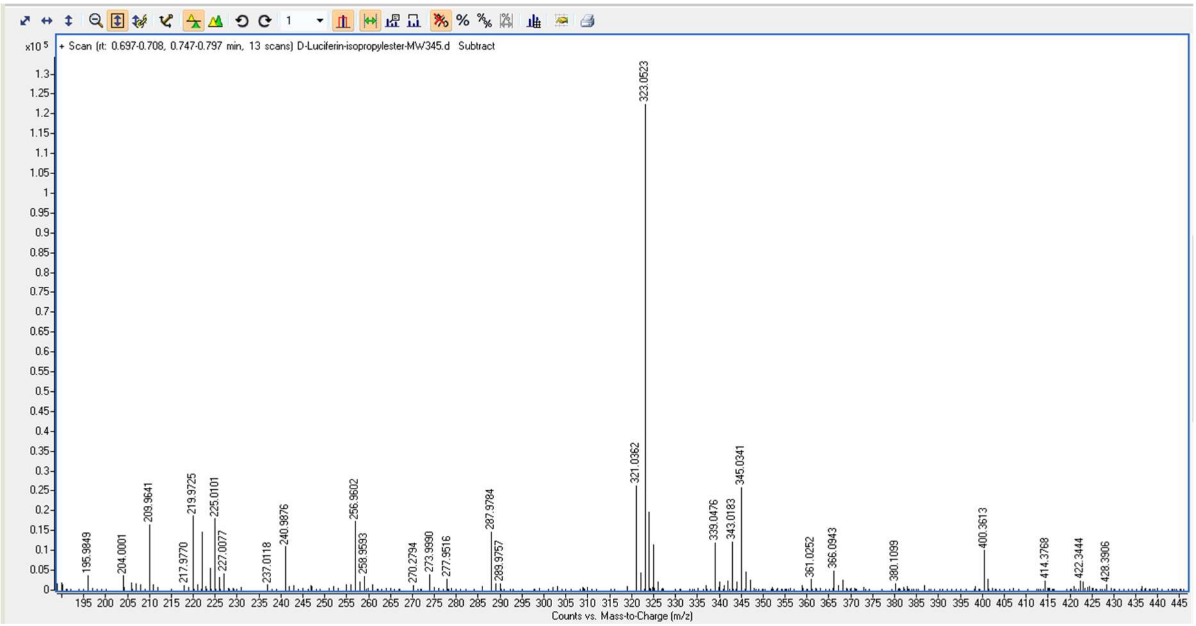


High Resolution Mass Spectrum for D-luc-*i*Pr (expected m/z 323.0519 for $[M+H]^+$).

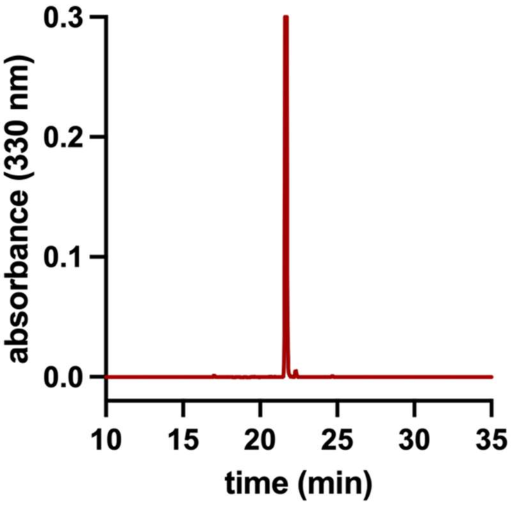


Analytical HPLC Chromatogram of D-luc-*i*Pr.

**D-luc-*n*Bu**

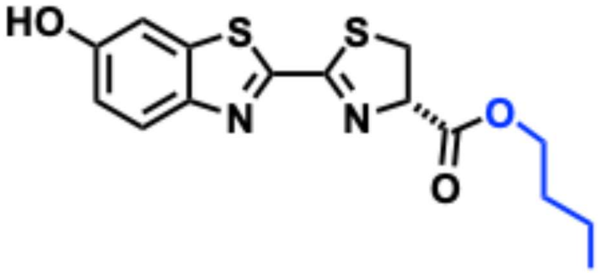

**Molecular formula:** $C_{15}H_{16}N_2O_3S_2$. **HRMS:** *m/z* calcd for $[M+H]^+$, 337.0675; found, 337.0680. **$^1$H NMR** (DMSO-$d_6$) δ 10.19 (s, 1H, OH), 7.95 (d, 1H, ArH), 7.43 (s, 1H, ArH), 7.06 (d, 1H, ArH), 5.48 (t, 1H, N–CH), 4.16 (t, 2H, O–$CH_2$), 3.79 (dd, 1H, S–$CH_2$), 3.67 (dd, 1H, S–$CH_2$), 1.56 (m, 2H, $CH_2$), 1.33 (m, 2H, $CH_2$), 0.89 (t, 3H, $CH_3$). **$^{13}$C NMR** (DMSO-$d_6$) δ 169.9, 165.0, 157.4, 156.6, 146.2, 137.2, 124.9, 117.2, 106.8, 77.8, 64.9, 34.6, 30.1, 18.7, 13.5.

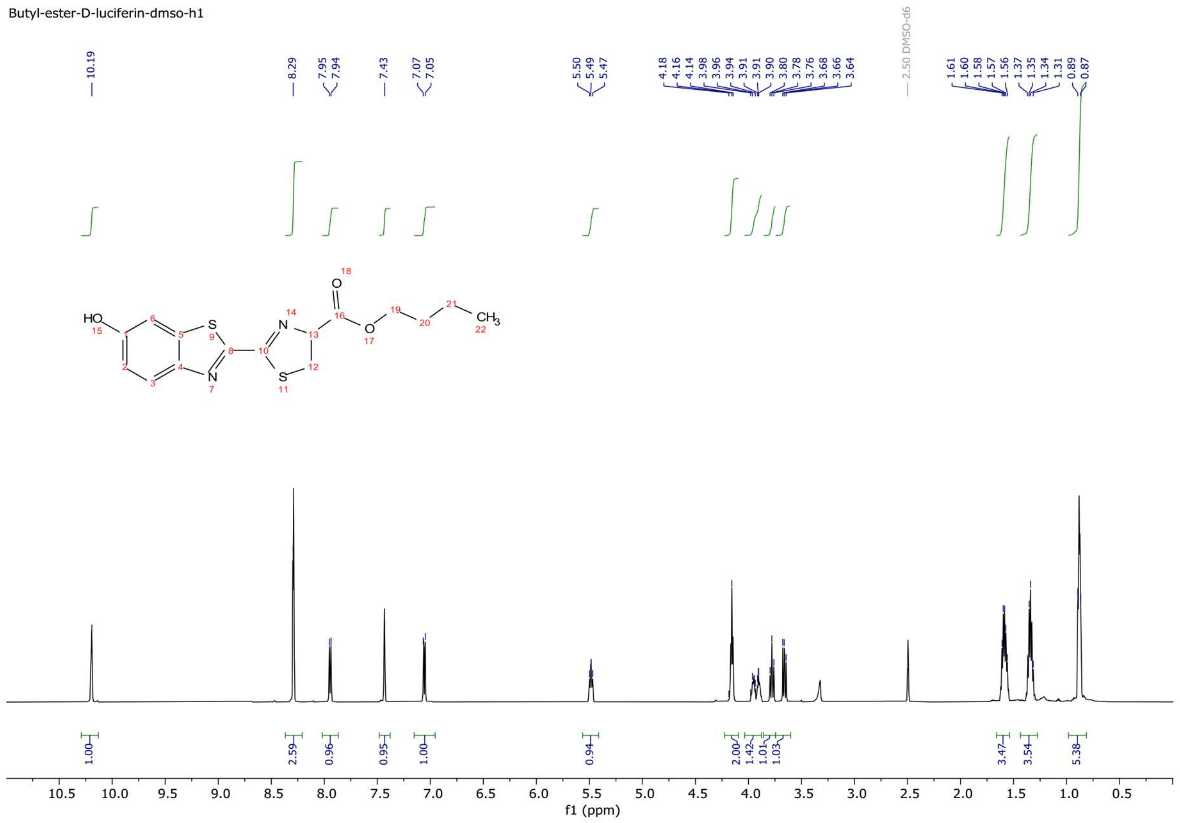

$^{1}H$ NMR spectrum of D-luc-*n*Bu.

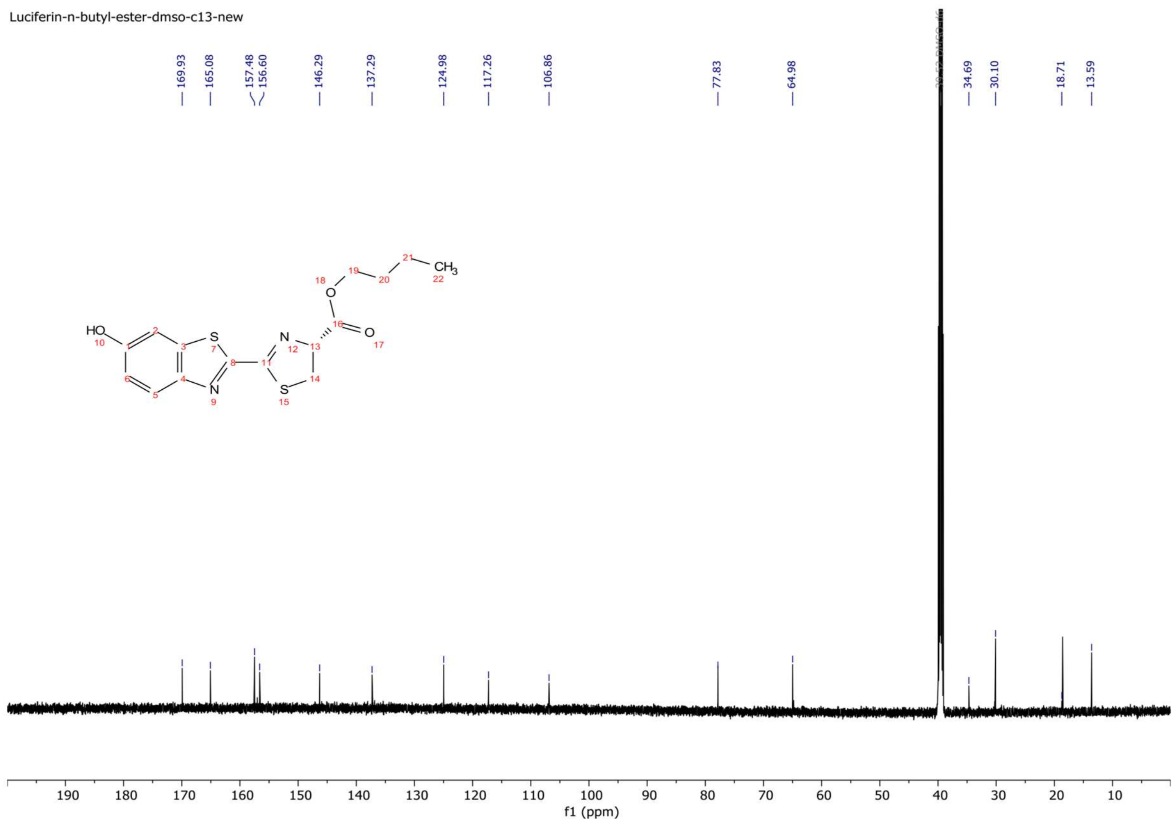


$^{13}C$ NMR spectrum of D-luc-*n*Bu.

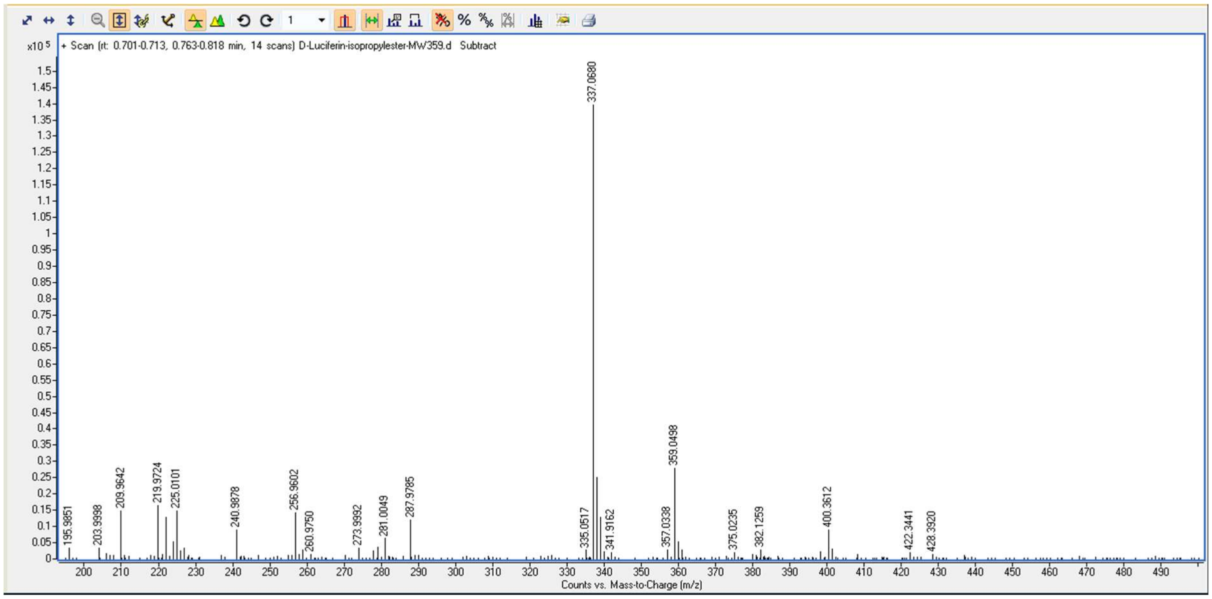


High Resolution Mass Spectrum for D-luc-*n*Bu (expected m/z 337.068 for $[M+H]^{+}$).

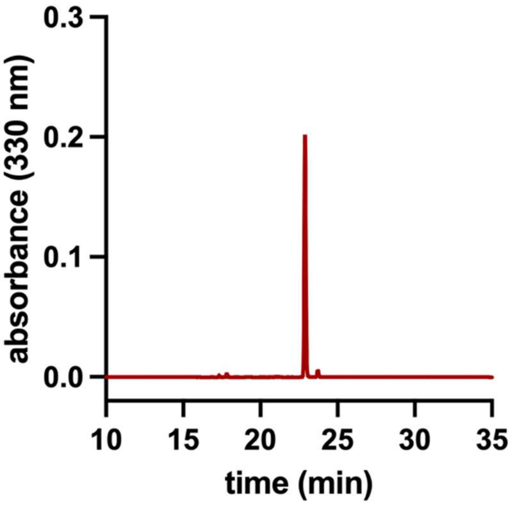

Analytical HPLC Chromatogram of D-luc-*n*Bu.

**D-luc-tBu**

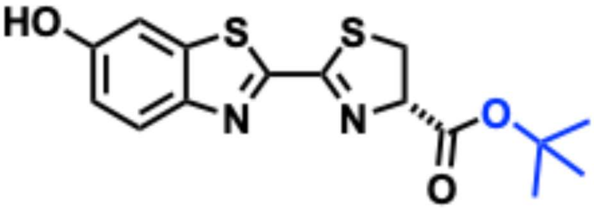

**Molecular formula:** $C_{15}H_{16}N_2O_3S_2$. **HRMS:** *m/z* calcd for $[M+H]^+$, 337.0675; found, 337.0677. **$^1$H NMR** (DMSO-$d_6$) δ 10.21 (s, 1H, OH), 7.95 (d, 1H, ArH), 7.44 (s, 1H, ArH), 7.07 (d, 1H, ArH), 5.37 (t, 1H, N–CH), 3.76 (dd, 1H, S–$CH_2$), 3.61 (dd, 1H, S–$CH_2$), 1.47 (s, 9H, $C(CH_3)_3$).

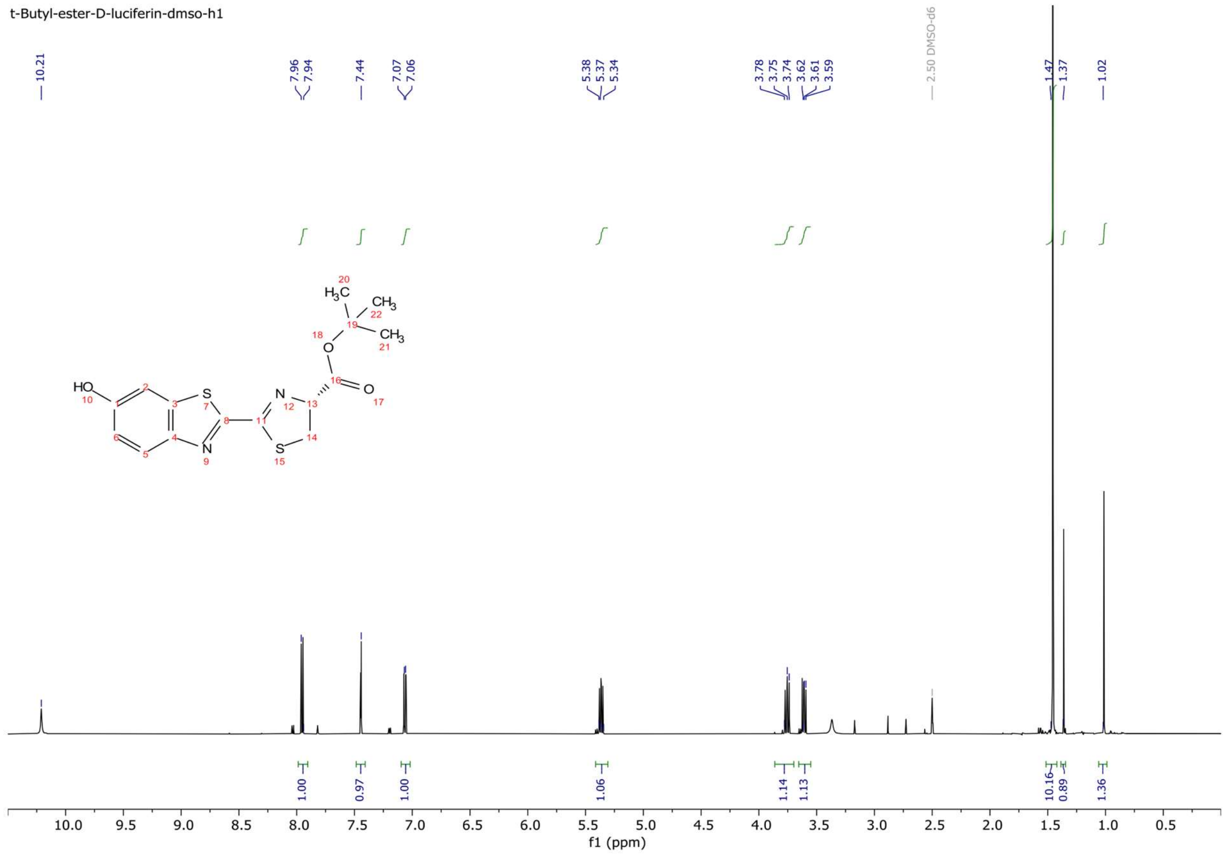


$^1$H NMR spectrum of D-luc-*t*Bu.

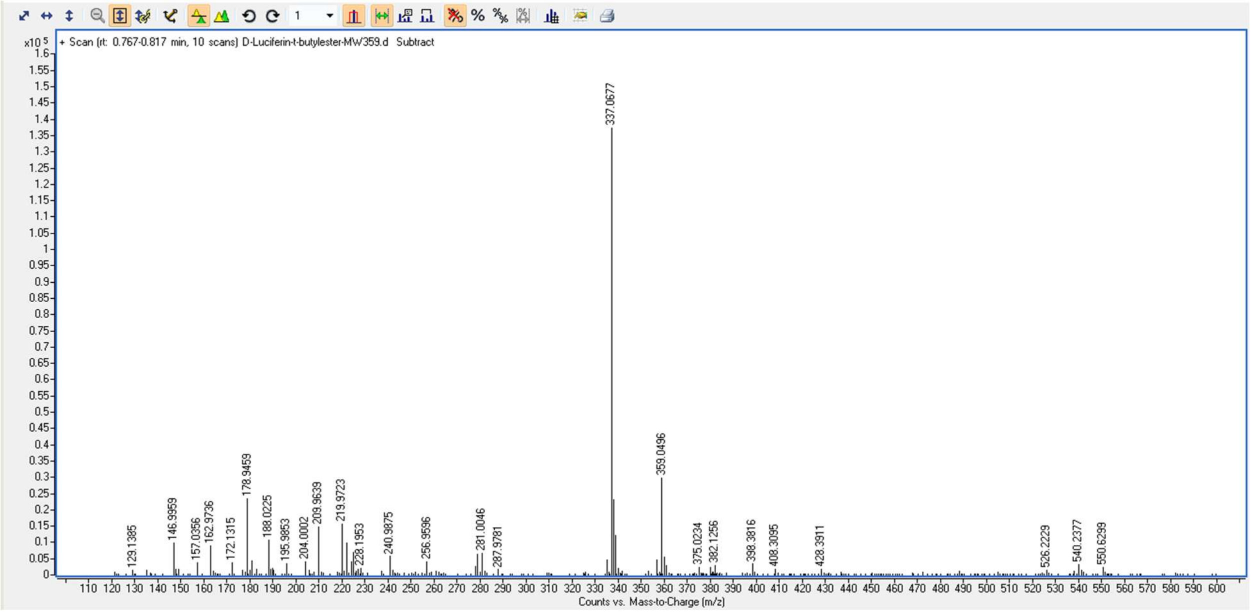


High Resolution Mass Spectrum for D-luc-*t*Bu (expected m/z 337.0675 for $[M+H]^+$).

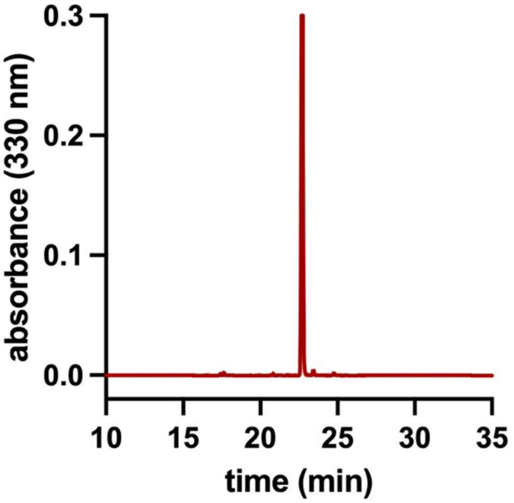


Analytical HPLC Chromatogram of D-luc-*t*Bu.
